# Anticipatory energization revealed by pupil and brain activity guides human effort-based decision making

**DOI:** 10.1101/2020.02.25.964676

**Authors:** Irma T. Kurniawan, Marcus Grueschow, Christian C. Ruff

## Abstract

An organism’s fitness is determined by how it chooses to adapt effort in response to challenges. Actual effort exertion correlates with activity in dorsomedial prefrontal cortex (dmPFC) and noradrenergic pupil dilation, but little is known about how these neurophysiological processes guide decisions about future efforts: They may either provide anticipatory energization helping to accept the challenge, or a cost representation weighted against expected rewards. Here we provide evidence for the former, by measuring pupil and fMRI brain responses while humans chose whether to exert efforts to obtain rewards. Pupil-dilation rate and dMPFC fMRI activity related to anticipated effort level, with stronger correlations when participants chose to accept the challenge. These choice-dependent effort representations were stronger in participants whose behavioral choices were more sensitive to effort. Our results identify a process involving the peripheral and central human nervous system that guides decisions to exert effort by simulating the required energization.

## Introduction

Should I go to the gym tonight or should I skip training? Such trade-offs between effort and reward are commonplace in our everyday lives. In fact, the ability to choose between high cost, high yield or low cost, low yield actions is crucial for survival in all animals (Bautista, Tinbergen, and Kacelnik 2001). Reward signals found in the dopaminergic (DA) and core brain reward circuitry have long been identified to play a pivotal role in appetitive motivation and in guiding choices (Schultz, Dayan, and Montague 1997; Bartra, McGuire, and Kable 2013; Niv, Daw, and Dayan 2005; Beierholm et al. 2013; Varazzani et al. 2015; Schultz 2002; Walton and Bouret 2019; Ostlund et al. 2011). By contrast, it is much less clear how decisions may be guided by effort signals. Previous work has indicated that neural signals for effort in the noradrenergic (NA) neuromodulatory arousal system (Varazzani et al. 2015; Zénon, Sidibé, and Olivier 2014) and fronto-insular network (Aridan et al. 2019; Arulpragasam et al. 2018; Kurniawan et al. 2013; Skvortsova, Palminteri, and Pessiglione 2014; Hauser, Eldar, and Dolan 2017; Meyniel et al. 2013; Prevost et al. 2010) scale monotonically with increasing task-difficulty levels (McGuire and Botvinick 2010), but how these neuromodulatory processes and neural representations functionally contribute to the choice process and goal-directed behaviour is unknown.

Two possible functional roles of effort signals have been proposed. First, a prevailing view in decision theory posits that efforts incur action ***costs*** that are weighed against the rewards to compute the net value of the action (Hull 1943). Consistent with this view, several human functional magnetic resonance imaging (fMRI) studies show net value signals for reward that are subjectively “discounted” by effort (Aridan et al. 2019; Arulpragasam et al. 2018; Bernacer et al. 2019; Chong et al. 2017; Prevost et al. 2010; Burke et al. 2013; Klein-Flügge et al. 2016). However, these net value signals primarily reflect the rewarding aspects of the choice options, which impairs direct interpretations whether these signals truly reflect effort and how effort per se may impacts on the choice process.

Second, consistent with the idea that effort represents resource mobilization (Hockey, G. Robert 1997), decisions may require an estimation of the ****energization**** needed to ensure that the action under consideration can be successfully achieved (Paravlic et al. 2018). A sizeable literature indicates that locus coeruleus noradrenergic (LC-NA) activity plays an important role in changing arousal states (Pfaff, Martin, and Faber 2012; Takahashi et al. 2010; Poe et al. 2020) by providing neuromodulatory input to the entire neocortex (Porrino and Goldman‐Rakic 1982; Chandler, Lamperski, and Waterhouse 2013; Schwarz et al. 2015), thereby facilitating energization (Varazzani et al. 2015; Jahn et al. 2018). NA activity can directly influence pupil size and is tightly linked to changes in pupil dilation (Joshi et al. 2016; Reimer et al. 2016; Gelbard-Sagiv et al. 2018), making phasic, task-related pupil an accurate indicator of brain arousal states (Yüzgeç et al. 2018; McGinley et al. 2015). However, it remains unclear whether the effort signals that guide choices would also draw on the same pupil-linked NA arousal system that has been found to facilitate actual behavior energization (Varazzani et al. 2015; Zénon, Sidibé, and Olivier 2014; Borderies et al. 2020; Xiang et al. 2019).

Teasing apart these two scenarios is not trivial. One effective way forward is to investigate how signals that scale with effort levels differ depending on choice outcomes. Namely either “Yes” decisions, in which we choose to engage effort (e.g., exercising at the gym) versus “No” decisions whereby we forego the effort (Kurniawan et al. 2010). In a cost scenario, stronger brain signals for effort (after controlling for rewards) would decrease the option’s net value and push individuals towards a “No” decision. Thus, a cost scenario would predict a steeper neural effort signal in “No” compared to “Yes” decisions. In an energization scenario, by contrast, a higher effort signal would trigger readiness to mobilize resources and tip individuals towards a “Yes” decision. The energization scenario would therefore predict the opposite pattern of steeper effort-related signals during “Yes” compared to “No” decisions (Fig. 1A).

**Figure 1.**
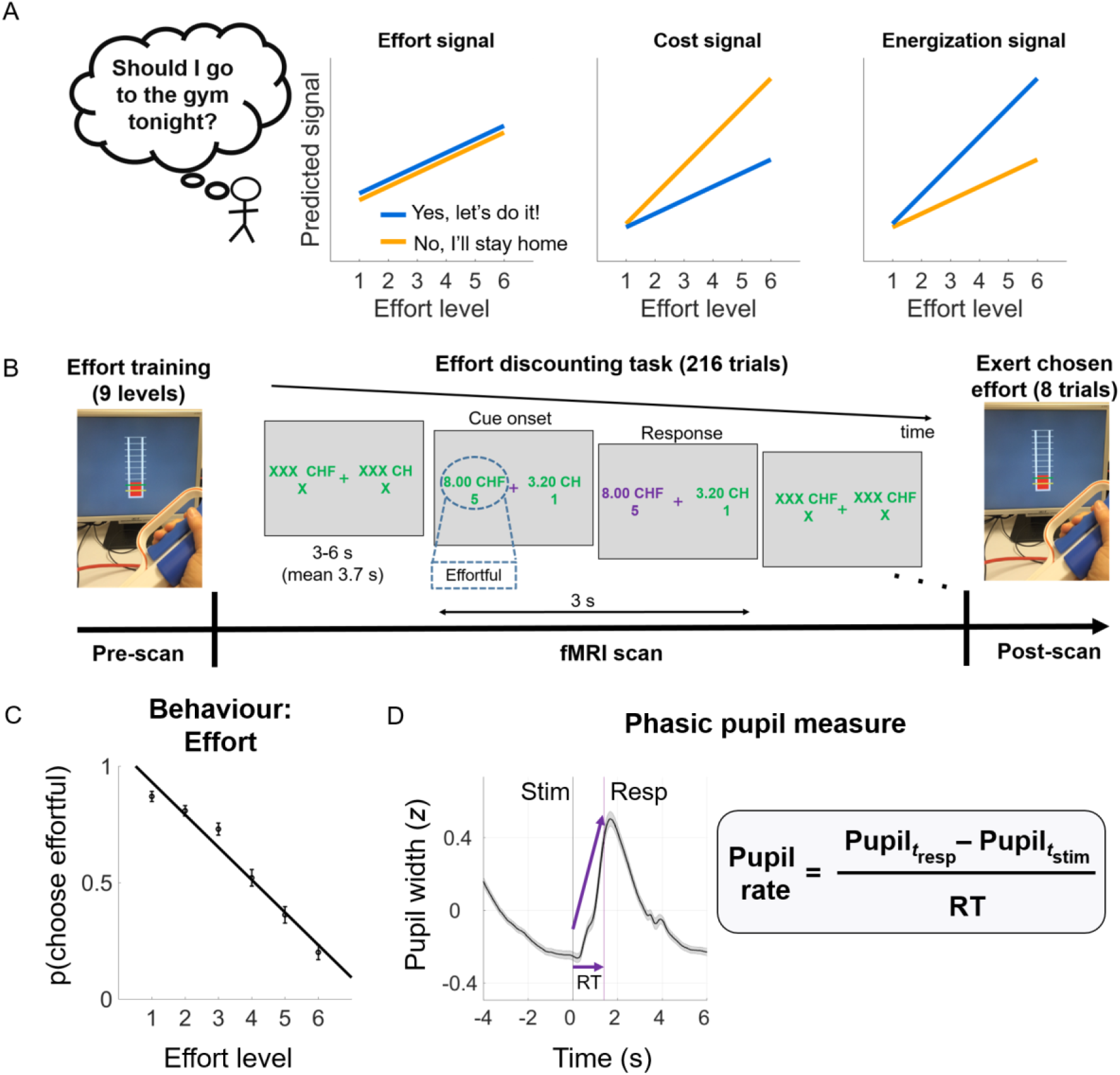
Predictions, task design, and key measures. **A) Three possible patterns of anticipatory neural responses to effort.** Left: Signals coding for effort per se would scale monotonically with effort regardless of choice. Middle: Signals coding for the decision cost associated with effort should be steeper across effort levels when individuals reject the effort. Right: Signals coding the anticipatory energization needed to accept the challenge should be steeper across effort levels when when individuals accept the effort. **B) Experimental paradigm**. Pre-scan: Participants received visually-guided effort training on a hand-held dynamometer. Levels 1-9 correspond to 10-90% maximum voluntary contraction (MVC). In the fMRI scanner, participants chose between an effortful option associated with variable amounts of reward and effort and a non-effortful option with smaller reward. Post-scan: Outside the scanner, eight randomly selected trials were realized and participants executed the effort they chose to obtain the reward. **C) Behavioural effort sensitivity**. This individual measure was derived by calculating for each participant the slope of the probability to choose the effortful option across effort levels. **D) Phasic pupil measure**. Grand-mean of pupil width during decision making showed a stereotypical dilation shortly following stimulus onset, peaking right after averaged response onset (purple line), and constricting down to baseline level around stimulus offset. Pupil rate (z/s) was calculated by subtracting pupil width at response from pupil width at stimulus onset, divided by response times (RT).

Here we apply this experimental logic, using an effort/reward tradeoff task in an fMRI setting, while simultaneously tracking pupil dilation, a putative marker for LC-NA firing. This combination allows us to investigate systematically to what degree the brain arousal system may encode anticipated effort during decision making as a cost or energization signal.

First, we explored whether pupil-linked arousal, as measured in rate of pupil change (Joshi et al. 2016; Reimer et al. 2016), scales monotonically with increasing effort, and if such effort sensitivity in the pupil rate differs depending on choice outcome (“Yes” vs “No”).

Second, at the neural level, we similarly examined whether known cortical representations of effort reflect a neural version of such choice-dependent effort signal. Based on previous work with a similar paradigm (Kurniawan et al. 2013; Skvortsova, Palminteri, and Pessiglione 2014; Meyniel et al. 2013; Prevost et al. 2010; Hauser, Eldar, and Dolan 2017), we expected these signals to be localized within the fronto-insular network, which based on its connectivity to the LC (Poe et al. 2020) may be strongly affected by NA arousal processes.

Third, if such effort signaling is at all behaviorally relevant, then we expect individuals who show stronger choice-dependent effort signals in pupil and the brain to display stronger effort sensitivity in their behavior, namely in choice frequencies. In the cost scenario, we would expect behavioral effort sensitivity to be positively correlated with the difference in effort scaling of “No” > “Yes” decisions, since individuals who assign higher costs to effort should forego the effort challenge more often. The energization scenario, by contrast, would predict behavioral effort sensitivity to be positively correlated with the difference in effort scaling of “Yes” > “No” decisions, since those behaviorally more affected by effort would need a stronger energization signal to accept a given effort level.

Fourth, we conducted a series of control analyses to ascertain that the observed effects were not driven by changes in choice difficulty and reward value of the options. Moreover, since endogenous fluctuations of arousal states may cause a general bias towards exerting effort (Murphy, Vandekerckhove, and Nieuwenhuis 2014), and since elevated emotional arousal prior to a force-production task can increase voluntary effort (Schmidt et al. 2009), we also controlled for effects of tonic pupil signals as indexed by pre-trial pupil baseline level (PBL).

## Results

In the fMRI scanner, participants made a series of effort/reward tradeoff choices between an effortful option and a non-effortful option (Fig. 1B). On each trial, the effortful option entailed varying amounts of effort (1 of 6 levels, 40-90% maximum voluntary contraction—MVC; shown as levels 4-9) and reward (1 of 6 levels, 0.5-10 CHF; Fig. 2A). The non-effortful option entailed minimal effort (fixed at level 1) and a lower reward amount (30 or 40% of the reward amount of the effortful option). Each effort to be considered entailed 10 repetitions (‘reps’) of hand muscle contractions (3 s) and relaxations (3 s) and was implemented outside the scanner 30-60 minutes after the experiment. Indeed, during the scan participants were not provided with a hand dynamometer device and thus were fully aware that they would make successive decisions without executing the force task. We implemented this temporal separation between decisions and actual exertion to set up a hard test whether arousal effects could still be observed in cases where post-decisional motor preparation was completely absent. Given this experimental design, any phasic arousal effect could not be due to an impending motor action, and any lack of such an effect would unlikely be due to the effort task being hypothetical or trivial. We could thus investigate whether pupil-linked arousal scales with increasing physical effort during mere mental simulation when deciding about future efforts.

**Figure 2.**
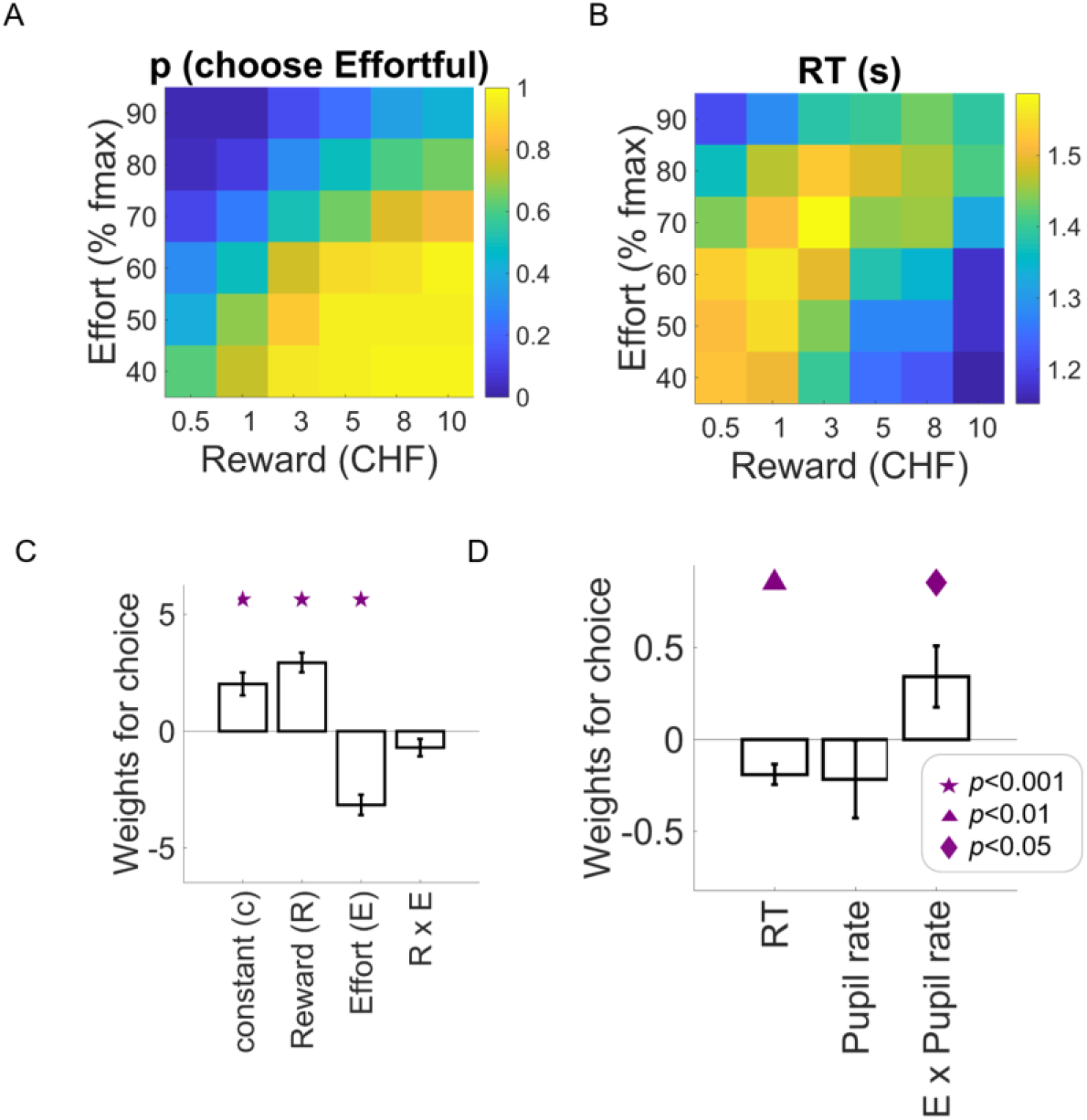
Behavioral and pupil results. Choice proportions (A) and RT (B) as a function of reward and effort associated with the effortful option. **C)** Weights of logistic regression of choice (1=effortful; 0 non-effortful) on reward, effort, and the interaction from a ‘standard’ model based on the offers the participants see on the screen. **D)** Weights of logistic regression of choice on RT, pupil rate, and effort-by-pupil rate interaction from an extended model (containing the standard model, RT, pupil rate, and other variables; see Supplementary Materials). This extended regression (D) had a higher model-fit (adjusted R-squared) than the standard one (C), t(48)=5.35, p<0.0001, suggesting that pupil measures together with other task parameters such as reward, effort, and RT, can explain choice above and beyond the ‘standard’ option attributes (reward and effort). Symbols indicate significance levels against zero. Bar plots display means ± 1 standard error of the mean (SEM).

### Systematic effort-reward trade-offs during choice

Initial analyses confirmed that participants indeed systematically traded off the proposed efforts and rewards when making decisions (Fig. 2A), as expected based on previous work (Prevost et al. 2010; Kurniawan et al. 2010; Chong et al. 2017). Effortful options were selected significantly more often when they offered higher rewards and lower effort amounts (Fig. 2C; logistic regression of choice; 1=choose effortful, 0=choose non-effortful; *N*=49; adjusted R^2^ *M*=0.62, *SEM*=0.017; *t*_reward_(48)=6.93, *p*<0.0001; *t*_ef f ort_(48)=-7.25, *p*<0.0001). In particular, effortful options were selected / abandoned most often when they were clearly attractive (high rewards for low effort) / unattractive (low rewards for high efforts), although the interaction effect was only marginally significant (*t*_reward*ef f ort_(48)=-1.93, *p*=0.06). This ‘standard’ logistic regression model confirms previous findings that decisions vary as a function of the offered rewards and the required effort. Furthermore, we found evidence in response times (RT) data (Fig. 2B) that choice outcome may further reveal information about the decision process. Multiple regression of RT (z-scored) confirmed significant effects of reward and effort (*N*=49; adjusted R^2^ *M*=0.22, *SEM*=0.014; *t*_reward_(48)=3.93, *p*=0.0003; *t*_effort_(48)=-5.90, *p*<0.0001). In addition, RTs were faster when participants selected the effortful option than when they selected the non-effortful option (*t*_choice_ (48)=-4.46, *p*<0.0001; other effects: *t*_choice*reward_(48)=-5.82, *p*<0.0001; *t*_choice*ef f ort_(48)=8.44, *p*<0.0001; *t*_constant_(48)=6.68, *p*<0.0001; *t*_reward*ef f ort_(48)=-0.8, *p*=0.41; *t*_choice*reward*ef f ort_(48)=1.3, *p*=0.019).

### An energization signal in the rate of pupil change

We then investigated whether pupil change rate contained information correlated with choice outcome, over and above the known effects of reward and effort. To this end, we added pupil measures to the ‘standard’ logistic regression of choice (Fig 2C). This extended regression (Fig. 2D) replicated the effects of reward and effort, (*N*=49; adjusted R_2_ *M*=0.65, *SEM*=0.018; *t*_reward_(48)=6.56, *p*<0.0001; *t*_effort_(48)=-7.39, *p*<0.0001), and also revealed a significant reward-by-effort interaction, *t*_reward*ef f ort_(48)=-2.41, *p*=0.019. Crucially, the extended regression revealed a significant interaction between effort level and pupil rate, *t*_ef f ort*pupil_rate_(48)=2.04, *p*=0.04 (see supplementary materials for full statistics of the extended regression).

To examine whether this interaction effect reflects stronger effort representations for “yes” choices (i.e., energization) or for “no” choices (i.e., a cost signal, see Fig 1A), we directly examined the slopes of the regressions of pupil signals on anticipated effort levels during both types of choice outcomes. Averaged across both types of outcomes, the regression slope was indeed positive (one-sample t-test on averaged effort slopes across choice: *t*(48)=3.24, *p*=0.002) but importantly, it was significantly steeper when participants chose the effortful option compared to when they chose the non-effortful option, effort-by-choice interaction, *t*(48)=2.59, *p*=0.012 (Fig 3C). Thus, the pattern of effort representations in pupil signal during “yes” and “no” choices is consistent with the scenario that arousal system engagement during choice relates to energization for the future challenge that is being pondered.

**Figure 3.**
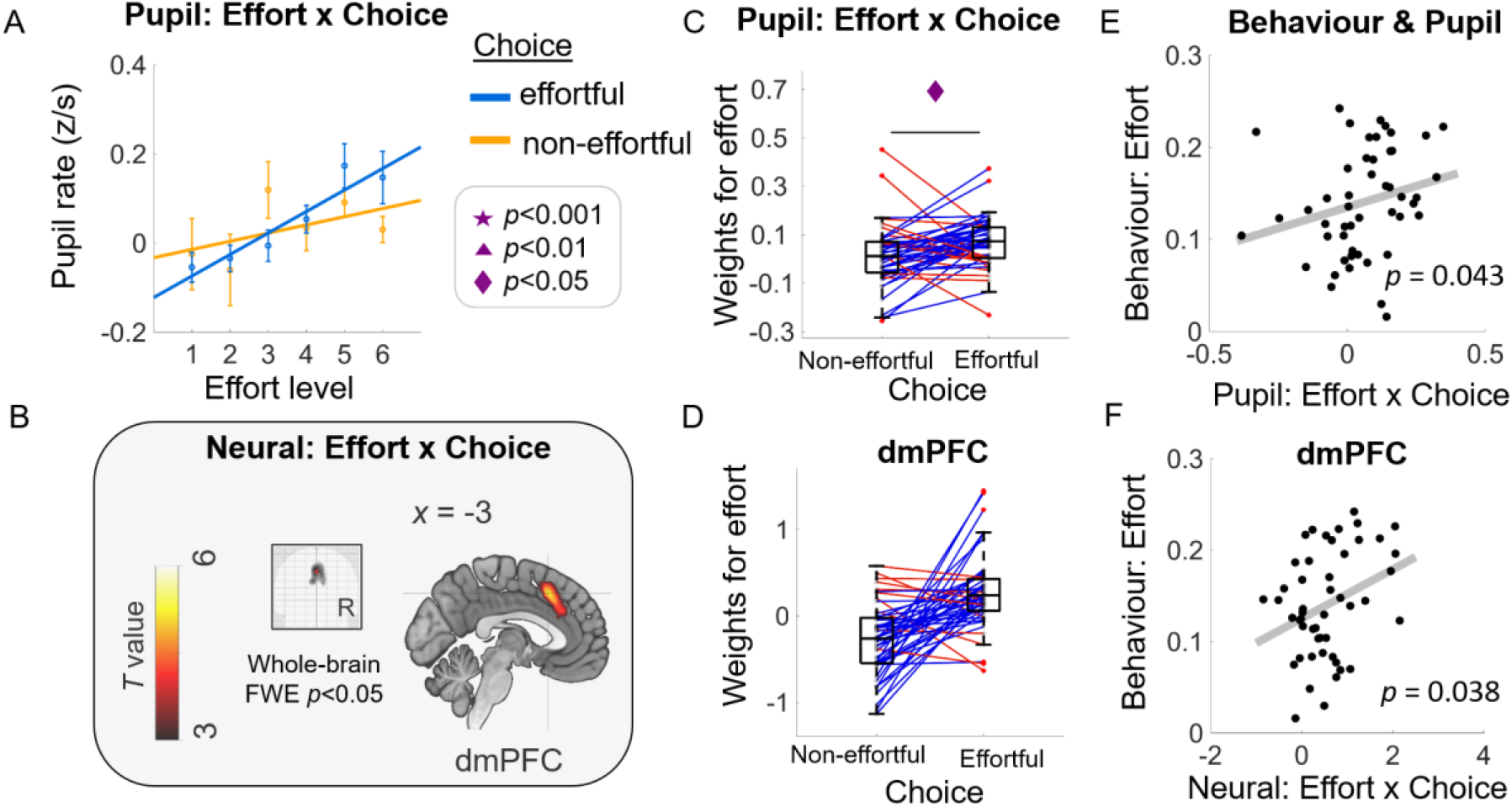
Energization signals in pupil and brain activity correlated with behavioural effort sensitivity. Consistent with the energization scenario, effort representations in pupil (A) and in the brain (B) are higher when participants accepted compared to rejected the effortful option (Choice “effortful” versus “non-effortful”). This significant effort-by-choice interaction effect is evidenced by higher effort beta weights when participants chose the effortful versus the non-effortful option in pupil (C) and in extracted BOLD signal change within dmPFC functional ROI (D; displayed solely for illustration purposes; no statistical test was done). Both the pupil (E) and neural (F) energization signals were positively correlated with individual behavioural measure of effort sensitivity as shown in Fig 1C. Panel A: Dots with error bars represent means ± 1 SEM. Lines are linear fits of the means (using the MATLAB polyfit(x,y,1) function). Panel B: Glass-brain image and sagittal slice showing that BOLD amplitude in dmPFC is uniquely correlated with effort-by-choice regressor. Panels C & D: Boxplots display the median (central line), 25^th^ and 75^th^ percentiles (bottom and top edges), and non-outlier low and high extreme values (bottom and top error bars). Blue lines show subjects whose effort slope is higher in effortful choice than in non-effortful choice, red lines show subjects who have the opposite effect. Panels E & F: Each data point represents a subject. P-values represent significance level from robust regressions.

### Neural responses in dmPFC also reflect energization

To identify neural processes that may similarly reflect energization, we then examined BOLD responses during the decision process. Analysis of the brain responses time-locked to the presentation of the options (stimulus onset) revealed a significant, and structurally similar, effort-by-choice interaction in dmPFC (covering both SMA and ACC; peak MNI space coordinates: [-3, 18, 45]; *t* value, 5.32; extent: 301 voxels; *p*< 0.0001 FWE; Fig. 3B; GLM1). No other brain areas showed signals that survived whole-brain FWE correction (Table 1). ROI analysis within the dMPFC functional cluster illustrates that the activity related to anticipated effort strength is indeed higher in trials where the effortful option was selected compared to foregone (Fig. 3D solely for illustration; GLM2). Thus, similar to the pupil signals described above, BOLD activity in dmPFC also shows anticipatory effort signaling in a way that is consistent with energization to overcome future physical challenges.

**Table 1.** MNI coordinates and statistics for GLM1: effort-by-choice, choice, reward, effort, pupil rate, and RT modulation. All effects are from t-tests. *P* values are at cluster-level FWE correction.

| Effect | Brain region | k | t-value | p-value | MNI Coordinates |  |  |
| --- | --- | --- | --- | --- | --- | --- | --- |
|  |  |  |  |  | x | y | z |
| Effort-by-choice (positive modulation) | L Superior Medial Gyrus | 301 | 5.324 | <0.0001 | -3 | 18 | 45 |
|  | L ACC |  | 4.786 |  | -6 | 27 | 27 |
| Effort-by-choice (negative modulation) | L Postcentral Gyrus | 4085 | 6.382 | <0.0001 | -33 | -42 | 57 |
|  | R Superior Frontal Gyrus |  | 6.248 |  | 24 | -6 | 66 |
|  | R Postcentral Gyrus |  | 6.242 |  | 30 | -42 | 57 |
|  | L Middle Temporal Gyrus | 147 | 4.688 | 0.009 | -60 | -30 | -3 |
|  | L Middle Temporal Gyrus |  | 4.151 |  | -57 | -51 | -6 |
| Choice (Effortful > non-effortful) | L Middle Frontal Gyrus | 146 | 4.749 | 0.02 | -30 | 30 | 36 |
| Choice (Non-effortful > effortful) | no supra-threshold clusters |  |  |  |  |  |  |
| Reward (positive modulation) | L Caudate Nucleus | 1320 | 7.417 | <0.0001 | -9 | 9 | 0 |
|  | R IFG (p. Orbitalis) |  | 6.752 |  | 36 | 21 | -9 |
|  | L IFG (p. Orbitalis) |  | 5.662 |  | -30 | 24 | -3 |
|  | R Middle Frontal Gyrus | 464 | 7.237 | <0.0001 | 39 | 21 | 27 |
|  | L Inferior Parietal Lobule | 1046 | 6.61 | <0.0001 | -36 | -63 | 51 |
|  | L Precuneus |  | 4.949 |  | -3 | -66 | 42 |
|  | L Middle Occipital Gyrus |  | 4.948 |  | -30 | -78 | 27 |
|  | R Middle Temporal Gyrus | 211 | 5.767 | 0.003 | 60 | -30 | -6 |
|  | R Middle Temporal Gyrus |  | 3.781 |  | 57 | -9 | -18 |
|  | R Fusiform Gyrus | 900 | 5.57 | <0.0001 | 24 | -81 | -9 |
|  | L Fusiform Gyrus |  | 5.044 |  | -24 | -78 | -9 |
|  | L Cerebellum (Crus 2) |  | 5.018 |  | -12 | -81 | -27 |
|  | L Middle Temporal Gyrus | 149 | 5.489 | 0.012 | -60 | -21 | -15 |
|  | L Superior Frontal Gyrus | 478 | 5.06 | <0.0001 | -21 | 36 | 48 |
|  | L IFG (p. Triangularis) |  | 4.857 |  | -39 | 21 | 24 |
|  | L Middle Frontal Gyrus |  | 4.756 |  | -36 | 12 | 57 |
|  | L Superior Orbital Gyrus | 115 | 5.021 | 0.03 | -30 | 51 | 3 |
|  | R Inferior Parietal Lobule | 597 | 4.922 | <0.0001 | 33 | -72 | 24 |
|  | R Inferior Parietal Lobule |  | 4.839 |  | 39 | -60 | 48 |
|  | R Middle Temporal Gyrus |  | 4.706 |  | 54 | -48 | 12 |
|  | L ACC | 167 | 4.484 | 0.007 | -6 | 42 | 12 |
| Reward (negative modulation) | no supra-threshold clusters |  |  |  |  |  |  |
| Effort (positive modulation) | R Rolandic Operculum | 151 | 5.503 | 0.008 | 48 | 3 | 12 |
|  | L Linual Gyrus | 365 | 4.482 | <0.0001 | -15 | -84 | 3 |
|  | L Middle Occipital Gyrus |  | 4.352 |  | -30 | -90 | 15 |
|  | L Calcarine Gyrus |  | 4.073 |  | 3 | -75 | 15 |
|  | R Calcarine Gyrus | 117 | 4.313 | 0.022 | 30 | -69 | 15 |
| Effort (negative modulation) | no supra-threshold clusters |  |  |  |  |  |  |
| Pupil rate (positive modulation) | R Precuneus | 8334 | 7.993 | <0.0001 | 6 | -78 | 42 |
|  | L Calcarine Gyrus |  | 7.485 |  | -3 | -90 | 0 |
|  | R Superior Frontal Gyrus | 1081 | 6.774 | <0.0001 | 21 | -9 | 69 |
|  | R Middle Frontal Gyrus |  | 6.639 |  | 45 | -9 | 57 |
|  | L Posterior-Medial Frontal |  | 6.041 |  | -18 | -12 | 72 |
|  | R Temporal Pole | 307 | 6.500 | <0.0001 | 54 | 15 | -6 |
|  | R Insula Lobe |  | 4.806 |  | 36 | 9 | 12 |
|  | L Rectal Gyrus | 287 | 5.715 | <0.0001 | -15 | 12 | -9 |
|  | R Caudate Nucleus |  | 4.519 |  | 9 | 6 | 3 |
|  | R Olfactory cortex |  | 4.220 |  | 21 | 9 | -15 |
|  | L Temporal Pole | 225 | 4.854 | 0.001 | -57 | 9 | -3 |
|  | L IFG (p. Opercularis) |  | 4.283 |  | -42 | 9 | 15 |
|  | L IFG (p. Orbitalis) |  | 3.563 |  | -30 | 27 | -3 |
|  | R Superior Temporal Gyrus | 138 | 4.600 | 0.013 | 57 | -39 | 27 |
| Pupil rate (negative modulation) | L Middle Temporal Gyrus | 123 | 5.914 | 0.02 | -51 | -36 | -3 |
|  | L Inferior Temporal Gyrus |  | 3.581 |  | -60 | -12 | -21 |
|  | L Angular Gyrus | 196 | 5.628 | 0.003 | -54 | -63 | 36 |
|  | L Superior Medial Gyrus | 280 | 5.168 | <0.0001 | -6 | 48 | 45 |
|  | R Superior Medial Gyrus |  | 4.652 |  | 3 | 30 | 60 |
|  | L Middle Frontal Gyrus | 96 | 4.703 | 0.044 | -33 | 24 | 51 |
| RT (positive modulation) | L Cuneus | 273 | 7.335 | 0.001 | 3 | -93 | 24 |
|  | R Cuneus |  | 6.818 |  | 6 | -84 | 42 |
|  | R Paracentral Lobule |  | 6.657 |  | 6 | -48 | 75 |
|  | R IFG (p. Orbitalis) | 219 | 5.795 | 0.003 | 45 | 36 | 0 |
| RT (negative modulation) | R Fusiform Gyrus | 11871 | -<br>14.006 | <0.0001 | 36 | -75 | -9 |
|  | L Fusiform Gyrus |  | -<br>12.751 |  | -36 | -69 | -9 |
|  | L Middle Occipital Gyrus |  | -<br>11.346 |  | -36 | -87 | 6 |

### Energization signals in pupil and dmPFC relate to behavioral effort sensitivity

To investigate whether the energization signals in pupil and dMPFC activity are indeed behaviorally relevant, we tested whether the difference in effort coding (slope across effort levels) between “yes” and “no” responses was associated with individual differences in how the anticipated degree of effort affected choice outcomes. For this analysis, we performed for each individual a simple logistic regression of choice on the associated effort levels (transformed such that a positive slope means higher likelihood to forego the option with increasing effort). The individual slopes of these regressions - our behavioral measure of effort sensitivity – were indeed positively correlated with the strength of each individual’s effort-by-choice effect in both pupil rate and dmPFC activity (taken from the ROI analysis), *robust regressions b*_pupil_rate_(47)=0.70, *p*=0.043; *b*_dmPFC_(47)=3.56, *p*=0.038; Fig. 3E-F). Thus, subjects with higher effort sensitivity (whose overall choice was more strongly affected by increasing effort) indeed showed, in both pupil rate and dMPFC activity, steeper effort coding when the effortful option was selected compared to when it was foregone. Therefore, the energization responses in pupil rate and the brain indeed appear to be relevant for guiding choices.

### Energization effects in pupil rate are independent of reward value, decision difficulty, or tonic arousal

It is theoretically possible that the effects we observed in pupil rate were driven by differences in reward or difficulty level of trials where effort is accepted versus rejected. Indeed, increases in pupil size have been observed for rewarding stimuli (Schneider et al. 2018) and trials that require greater cognitive control (van der Wel and van Steenbergen 2018). These effects might be confounded with the energization effect we reported, particularly because in some cases, high effort trials may be associated with high rewards, hence making the decision to either select or forego the effortful option more difficult. Our behavioral results had already contradicted these alternative explanations, since they were derived with statistical models that had accounted for any variance associated with reward levels and RT (an indirect proxy for decision difficulty (Kiani, Corthell, and Shadlen 2014)) (see Supplementary Materials). Nevertheless, to show more directly that the energization effect is clearly independent of reward and difficulty, we repeated the pupil analyses depicted in Figure 3 but now on the residuals of pupil rate after partialing out the effects of rewards and of RT (orthogonalization of pupil rate relative to these variables, one at a time). Once again, these control analyses revealed the effects already shown in top row of Figure 3, namely (1) stronger effort signals in residual pupil rate when participants accepted versus rejected the effortful option; *t*_resid_reward_(48)=2.59, *p*=0.012; *t*_resid_RT_ (48)=2.53, *p*=0.014 and (2) significantly positive associations between the pupil energization effect (effortful>non-effortful) and the behavioral effort-sensitivity parameter (*robust regression b*_resid_reward_(47)=0.71, *p*=0.043; *b*_resid_RT_(47)=0.68, *p*=0.048; (Fig. S4). Furthermore, to rule out an alternative explanation that the pupil is merely coding for any option attribute that participants experienced as result of their choice (in our case the other option attribute was reward), we replaced these analyses with a reward-by-choice interaction (instead of effort-by-choice). These control analyses yielded no significant reward-by-choice effects in the pupil data or the correlation with behavioral measure of reward sensitivity (Supplementary Materials, Fig S5). Thus, the energization effect we identified in pupil rate is independent of reward value, decision difficulty, or a reward-by-choice interaction, and thus reflects different neural mechanisms to those underlying conflict-driven pupil dilations and behavioral adjustments (Ebitz and Platt 2015).

To ascertain that our novel effect is also independent of ongoing background arousal, we defined the average pupil diameter during 500 ms prior to the presentation of the options as an index of pre-trial pupil baseline level (PBL). We did not find a relationship between PBL and choice frequencies (Supplementary material; Fig S2-3). This absence of a link between PBL and effort-based choice did not reflect more complex interactions with other experimental factors or influences from the previous trial, as ascertained by logistic regressions of choice on PBL, RT, reward, effort, and the interactions. Direct test of effort-by-choice interaction effect on PBL also yielded non-significant results (Supplementary Materials; Fig S2-3). Taken together, we thus found no evidence that ongoing background arousal state biases subjects to accept high-effort options, thus confirming the specificity of the energization effect for phasic arousal responses during the choice process.

### Energization effects in dMPFC are independent of neural representations of other task parameters

Importantly, we made sure that the observed energization effects in the dmPFC are indeed novel and separate from known neural correlates of effort-based decisions. To this end we had included the effort-by-choice interaction as a regressor together with main effects of choice, reward, effort, pupil rate, and RT in the same model without any orthogonalization (see methods). This allowed us to identify neural representations that are unique to each of the task parameters, ensuring that the effort-by-choice interaction cannot be explained by any combination of the other factors and allowing us to inspect our data for several known neural representations active during effort-based decisions (Fig 4).

**Figure 4.**
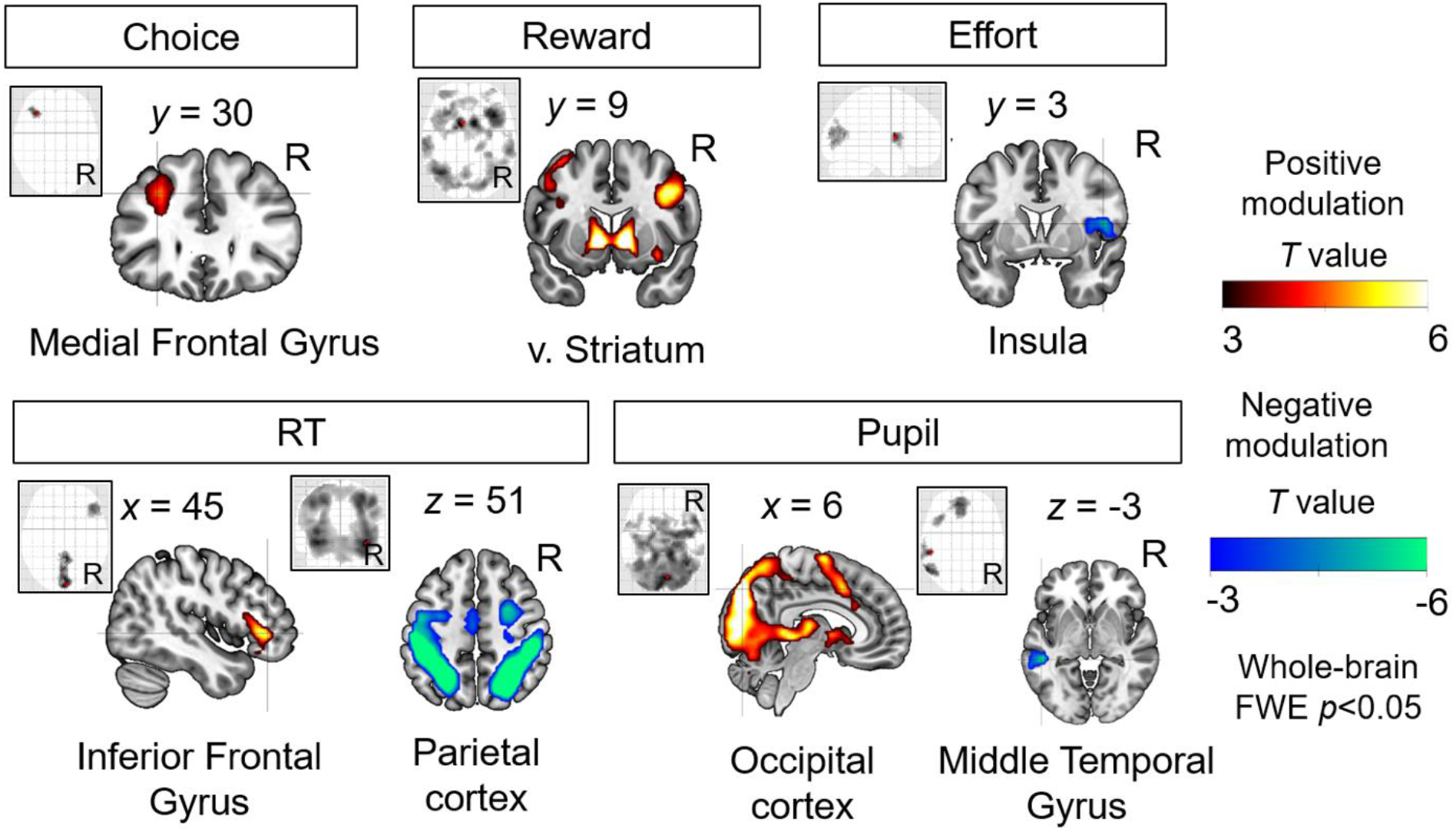
Neural representations of choice, reward, effort, RT, and pupil rate. These plots show whole-brain statistical parametric maps for neural representations of choice (effortful > non-effortful), reward, effort, RT, and pupil, p<0.05 FWE corrected. These established effects were derived with the same statistical model also used to identify the energization signals displayed in Figure 3; the latter signal is therefore specific and unrelated to these classic effects reported in the literature.

Consistent with previous demonstrations of the role of the dorsolateral prefrontal regions in executive function (Grueschow, Kleim, and Ruff 2020), we observed higher activity for choosing the effortful options compared to the non-effortful options in the left medial frontal gyrus. We also replicated previous findings of positive modulation of reward within the brain valuation system (Bartra, McGuire, and Kable 2013; Burke et al. 2013), with peak activity at the ventral striatum, and negative modulation of effort in the insula (Prevost et al. 2010). Moreover, we found slower button responses to be associated with higher activity in inferior frontal gyrus and faster responses to be associated with higher activity in a fronto-parietal network that is often implicated in task engagement (Dosenbach et al. 2008; Cole et al. 2013). Finally, we found faster pupil rate to be associated with lower amplitudes of BOLD responses to the presentation of the stimuli in the middle temporal gyrus. By contrast, faster pupil rate is associated with higher BOLD amplitudes in a large-scale network within the occipital cortex (extending to precuneus), consistent with established involvement of this network in visual processing (Goodale and Milner 2018). Thus, our brain results show that the energization signal in dmPFC is a conceptually new choice signal that is clearly distinct from previously observed effects of reward, effort, choice outcome, RT, and pupil signals (all reported effects survive whole-brain FWE correction, full statistics in Table 1). by reward value, decision difficulty, or background tonic arousal, thereby emphasizing the functional specificity of this energization signal.

Our results emphasize that phasic pupil-linked arousal during the decision process is tightly related to the amounts of effort that an individual agrees to invest, but they also raise the question what neural mechanisms may lie at the heart of this link between pupil and behavior. While the temporal sluggishness of the BOLD signal makes it difficult to provide a conclusive answer, we outline at least two plausible possibilities based on recent advances. First, simulating the required energization could trigger a “bottom-up” arousing influence that pushes decisions towards accepting effort. This would be consistent with the widely held view (Glimcher 2009) that the strength of neural representations for decision attributes directly influence choice – for instance, it has been shown that intensifying encoded rewards through simulation of future episodic events is linked with decisions that promote higher long-term pay-offs (Benoit, Gilbert, and Burgess 2011; Peters and Büchel 2010; Dassen et al. 2016; Bulley and Gullo 2017) and even with increases in prosocial behavior (Gaesser, Keeler, and Young 2018). Given this assumption, the arousal signal we observed in this study might either down-modulate anticipated effort costs or shift the decision rule (de Gee, Knapen, and Donner 2014), implying that a sufficiently strong arousal signal could bias a decision towards taking on the physical challenge. As for neural implementation, phasic LC activity is known to transmit feedforward information to ACC via ascending projections to prefrontal areas (Porrino and Goldman‐Rakic 1982; Schwarz et al. 2015; Chandler, Lamperski, and Waterhouse 2013), providing a plausible pathway for such bottom-up influences. Nervous readout of the autonomous arousal activation could provide a signal that the organism is indeed ready to take on the physical challenge, instantiating an additional mechanism to bias choices.

Second, simulated energization could simply be a byproduct of choice, implying a top-down influence from the cortical decision circuit to the arousal system. Decision outcomes could be relayed in the form of cortical descending input from the PFC into LC. ACC/dmPFC activity has been coupled with pupil diameter (de Gee et al. 2017; Ebitz and Platt 2015), and the timing of pupil modulation by ACC in some cases precedes that by LC (Joshi et al. 2016). Existing tracing data in rodents and monkeys also show afferent PFC projections as the main direct cortical influence on LC (Arnsten and Goldman-Rakic 1984; Dalsass et al. 1981). Intracranial stimulation in human ACC leads to subjective accounts of changes in arousal states, such as increased heart rate, coupled with the anticipation of challenges and a strong motivation to overcome difficult obstacles (Parvizi et al. 2013). This interpretation is also closely linked, though not identical, with the proposal that ACC computes the expected value of mobilizing mental resources (Shenhav, Cohen, and Botvinick 2016). Taken together, these observations are consistent with the idea of a top-down influence from dmPFC to the NA arousal system (Aston-Jones and Cohen 2005; Grueschow, Kleim, and Ruff 2020) that may serve to transmit information about the commitment to overcome great physical demand, thus resulting in speeded upregulation of arousal states to prepare the organism for the future challenge associated with the recent choice. Future studies may need to employ neuroimaging methods with higher temporal resolution to disambiguate these two hypotheses. Such studies may also employ pharmacological manipulation to increase NA tone activity, bio/neuro-feedback with pupil/LC activity, and mental simulation training (Steinmetz, Tausen, and Risen 2018) to increase arousal in a bottom-up fashion.

In our study, future efforts were signaled by the pupil-linked arousal system and dmPFC activity during choices that preceded actual exertion by about one hour. These results may seem at odds with those of monkey studies employing LC electrophysiology and NA pharmacology, which clearly showed effort sensitivity in the NA-system only during force production, but not during cues just moments prior to the effort (Jahn et al. 2018; Varazzani et al. 2015). The differences between our results and these datasets may reflect the very different time-periods separating choices from effort execution: In a paradigm such as ours, where the decisions pertain to efforts that have to be exerted sometime in the future (within 1 hour), the brain may need to perform a mental estimation of the amount of resources that will have to be mobilized in order to make the decision. This kind of simulation may not be needed, or may even be counterproductive, when decisions and exertions occur within seconds of one another. These methodological differences are not specific to our case but are rather a reflection of the state of the literature: Many monkey studies presented forced or choice cues that directly preceded actual exertions, whereas many human studies presented choice cues involving efforts that are delayed or even hypothetical. We clearly need studies that systematically investigate how the different timecourses present in these experiments affect effort coding in the NA arousal system and throughout the brain.

What would be the cognitive purpose of imagining or simulating behavior energization associated with a choice? Such simulation may contribute to metacognitive processes that evaluate the quality of our ongoing decisions to optimize future decision making (Fleming and Daw 2017). For an example from another domain, there is evidence that actual experience of choice and success in obtaining a food item influences how we value the food item in the future (Vinckier et al. 2018). Effort simulation may thus serve as a rich milieu for ‘scene construction’ (Hassabis and Maguire 2007) in which subjects evaluate the quality of their decision, which has the potential to shift future valuation. In our context, the source of simulation may include drawing from memory how much cognitive control needs to be mobilized (Shenhav, Cohen, and Botvinick 2016) in order to keep exerting physical effort rather than quitting, or retrieving the memory of previously incurred metabolic signal that accumulated the longer subjects exerted physical effort (Meyniel et al. 2013). Future experiments may directly test this conjecture by devising mental simulation paradigms in which participants imagine these specific elements of the force task, namely the sensations of mental fatigue or pain, and assessing how vividness ratings of these imagined bodily sensations would correlate with brain activity and choice. Furthermore, a mental simulation paradigm that manipulates agency might reveal stronger simulation signals for one’s own decisions compared to experimenter-imposed decisions, which would lend evidence for the use of simulation for self-evaluation (Fleming and Daw 2017).

Irrespective of these considerations, our results highlight that choices may be jointly guided by DA and NA systems for reward and effort processing, respectively. The majority of effort studies so far have reported a net value representation (reward discounted by effort) within the core brain valuation network (Prevost et al. 2010; Aridan et al. 2019) and in dmPFC (Klein-Flügge et al. 2016; Bernacer et al. 2019; Chong et al. 2017; Prevost et al. 2010; Burke et al. 2013; Arulpragasam et al. 2018). These fMRI results are consistent with animal data showing reduced willingness to choose a high-effort/high-reward option when dopamine is depleted (Salamone et al. 2007) and with the overarching dopaminergic role in motivational reward processing (Walton and Bouret 2019). Our present data concur with these previous studies, in showing reward coding within the brain valuation network (Prevost et al. 2010; Aridan et al. 2019) and notably NA-linked pupil dilations and dmPFC brain representations for physical effort (Kurniawan et al. 2013; Meyniel et al. 2013; Skvortsova, Palminteri, and Pessiglione 2014; Zénon, Sidibé, and Olivier 2014; Varazzani et al. 2015). This potential “partnership” of DA-coding for reward and NA-coding for effort does not seem to concur with the classical (but possibly simplistic) view that DA-linked reward processing is discounted in a subtractive fashion by NA-linked effort cost representations. We emphasize that our behavioral data and some aspects of our neural results are in line with previous computational suggestions that an option may be selected based on a trade-off between reward and effort (Fig 2). However, to our knowledge, prior work in humans has not examined how the effort sensitivity observed in the NA arousal system directly relates to choice. Here we were able to scrutinize this functional role using concurrent pupil-fMRI in an effort discounting task. Our results suggest that NA may play a complementary function to DA. Future studies may build on our results to further characterize the interaction between DA and NA, using the pupil rate measure in order to quantify energization signals that guide human decision making.

Variations in arousal states (measurable by pupil activity) - such as locomotion and sleeping - are coupled with oscillatory state changes in brain networks (Takahashi et al. 2010) that are thought to result from noradrenergic innervation to the cortex (Schwarz et al. 2015). However, there are also observations that arousal states may relate to movement during wakefulness and REM sleep, which are guided by cholinergic neuromodulatory projections from the basal forebrain to the cortex (Saper et al. 2010). This raises the concern whether we can truly draw the conclusions that our arousal effects evident in the pupil signals originate from LC-NA neuromodulation. While we cannot fully rule out potential effects of cholinergic activity in our study, a recent analysis with pupil activity and noradrenergic and cholinergic projections shed light on this issue, demonstrating that pupil rate in mice is more tightly linked with NA projections to the cortex, whereas activity in the cholinergic pathways more closely matched absolute pupil diameter (Reimer et al. 2016). Relatedly, a recent pharmacological study using clonidine to upregulate NA signaling in humans shows increased tonic pupil diameter during task-free intervals (Gelbard-Sagiv et al. 2018), but unfortunately does not report task-related phasic pupil rate, or a comparison with cholinergic signalling. Thus, data from mice generally support the view that our effects in pupil rate may reflect phasic arousal variations that most likely originated from NA-LC activity, but more investigation in humans are needed to replicate these findings.

Our results may have relevance for the diagnosis and therapy of brain disorders with deficits in motivated behavior. Committing to effort is a first step for success in motivated behaviors and the inability to commit to effort may bring about a cascade of clinical symptoms of apathy with a core feature of lack of self-initiated actions (Kurniawan, Guitart-Masip, and Dolan 2011; Husain and Roiser 2018; Le Heron, Apps., and Husain 2018). Recent neurocomputational work on effort-reward tradeoffs has identified promising phenotyping approaches of motivation disorders; these reflect key involvement of the fronto-subcortical circuitry and neuromodulatory systems including dopamine, serotonin, and noradrenaline (Meyniel et al. 2016; Pessiglione et al. 2018; Berwian et al. 2020). A specific role for noradrenaline is suggested by the finding that motivation deficits in depression that are inadequately treated by serotonergic antidepressants – including fatigue and loss of energy – have been shown to significantly improve following administration of NA (and dopaminergic) agents (Nutt et al. 2007). This highlights the critical yet overlooked role of NA in motivation regulation in depression (Moret and Briley 2011). Chronic exercise in mice also has been shown to increase LC-NA derived neuropeptide galanin that later conferred stress resilience (Tillage et al. 2020), providing further evidence of an adaptive role of NA-related energization signal. Our study contributes to this large body of work, by showing that the pupil-brain arousal system is sensitive to deliberations regarding sizable intensities of physical effort. Future work should further incorporate autonomic arousal and noradrenergic systems in quantitative models of motivation deficits (Pessiglione et al. 2018), particularly for dissociating arousal effects linked to anticipated effort from those that may reflect expected reward.

## Materials and Methods

### Participants

Fifty-two right-handed participants (29 females, mean age=22.3 (3) years) volunteered to participate in this study. We determined the sample size using power analysis based on the sma**l** to medium effect size (*d*=0.2-0.5) reported in past studies in the laboratory relating pupil size and biases in choice behavior (Raja Beharelle, et al., in prep; Grueschow et al., in prep). Participants received between 80-100 CHF (depending on the realized choices and performance) for their participation. Participants were screened for MRI compatibility, had no neurological or psychiatric disorders, and needed no visual correction. Data from one subject were excluded because of eye tracker data loss. Inclusion of this subject in the behavioral analysis did not change the statistical results, but for consistency, we excluded this data set from all analyses. We then screened subjects based on their mean choice proportion for the effortful option, p(choose effortful), to be within 0.1 and 0.9, and excluded data from one subject whose choice rate was 0.95. The final N was 49.

### Procedure

#### Force calibration

Upon arrival, participants were seated in the behavioral testing room, filled the MRI screening and consent forms, and received general instructions on the force task and MRI safety. Maximum voluntary force (MVC) level for each hand was obtained by averaging the top 33% force values produced during three 3-s squeezes. Continuous encouragement was given vocally during each entire squeeze period (e.g., “keep going, keep it up”).

#### Force training

Guided by a vertical bar on-screen (Fig. 1B), participants were trained to do hand squeeze sets at levels 10-90% MVC (displayed as levels 1-9). This dynamometer effort task mimics a typical hand force exercise at the gym, with a cycle of repetitions (‘reps’) of muscle contractions (3 s) and relaxations (3 s) for each level. To prevent muscle fatigue, these were done alternating between left and right hand. During training, one set consisted of 5 repetitions and there were in total 10 squeeze sets (10*5=50 reps) to be evaluated by a certain criterion. Levels 1-8 were presented once, pseudorandomly assigned to either left and right, and level 9 twice, once for each hand. The order of force levels was also pseudo-randomised. Half of the subjects practiced on levels 1, 3, 5, 7, 9 with left hand and 2, 4, 6, 8, 9 with right hand, and vice versa for the other half of subjects. The criterion was to maintain force above the target for at least two of the 3-s rep (non-consecutively). At the end of each training round, participants received a summary of their performance and were asked to repeat each unsuccessful force production. Overall, all participants underwent at most three training rounds (*M*=2.22, *SD*=0.46). After the last round, 38 participants successfully completed all 50 reps, whereas 11 participants had a few unsuccessful reps (*M*=4.3%, *SD*=3.5%). These results suggest that the training was very successful.

Following a 5-minute break, they proceeded with a subjective rating task in which they had to squeeze for each hand once at levels 1, 3, 5, and 9 for 5 s without knowing the difficulty levels. They were told that in some trials it would be easy to raise the bar to reach the target, which in this task was always displayed at the midline, while in other trials it would be harder to do it. After each 5-s squeeze, they then rated on a continuous visual analogue scale how effortful the grip was for them. They were instructed that the leftmost and rightmost point in the scale should refer to level 0 (merely holding the dynamometer) and level 10, respectively. The force training was successful as indicated by a close relationship between subjective and objective effort, mean pearson’s *r*=0.93, SEM=0.0073, *t*(46)=127.63, *p*<0.0001.

Prior to scanning, participants made five practice decisions and we made sure that participants fully comprehended the task. The effort discounting task was done in the fMRI scanner. Participants were aware that the effort they were considering now consisted of one set of 10 reps (instead of 5). To prevent participants from taking decisions based on anticipated muscle fatigue, only a random selection of eight decisions were actually realized in the behavioral testing room after the scan, and participants were fully aware of this. Participants then filled some questionnaires, were debriefed, given payment, and thanked for their participation before leaving the lab.

### Effort discounting task

In the scanner, participants were given a series of choices between an effortful and a non-effortful option. On each trial, the effortful option entailed varying effort (1 of 6 levels, levels 4-9) and reward amounts (1 of 6 levels, 0.5-10 CHF; Fig. 2A). The non-effortful option entailed minimal effort (fixed at level 1) and a lower reward amount (30 or 40% of the reward amount of the effortful option). To rule out risk as a potential confound (namely that accepting a level 9 offer gives a higher risk of task failure compared to accepting a level 4) we ensured that the effort training at all levels was successful (overall failure rate during training, *M*=0.9%, *SD*=2.4%),

We used a factorial design with six effort and six reward levels (36 cells) for the effortful option, and two reward levels for the non-effortful option. There were 3 trials in each cell, resulting in 6 x 6 x 2 x 3 = 216 trials. Trials were split in three fMRI runs of 72 trials (9 mins) and trial order was pseudorandomised per subject and run. The non-effortful option entailed effort fixed at Level 1 and smaller rewards (30-40% of the larger reward), giving a clear incentive to choose the non-effortful option if the larger effort was not worth the reward.

During a fixation period of 3-6 s (drawn from a gamma distribution with shape parameter 0.8 and scale parameter 1, mean 3.7s), the text indicating reward and effort levels was masked with a series of letters “X” (Fig. 1A). Following this period, the colour of the + sign at the centre changed and the effort and reward of each of the two options were presented on either side of the fixation point for a fixed duration of 3 s. This prompted the subjects that they were able to press either the left or the right key to indicate their choice. To provide decision feedback, this key response was promptly followed by a change in colour for the selected option. Regardless of key press, the stimuli remained on-screen for 3 s before the next fixation period was presented. If participants failed to respond during this period, the trial was coded as missing and no reward was gained. Amongst 49 participants, 13 had 1 missing trial, 5 had 2-5 missing trials, and 1 had 34 (15%) missing trials. Exclusion of this last subject did not change any result, so we decided to include them.

### Pupillometry

Participants’ right or left eye (depending on feasibility) was monitored using MR-compatible infrared EYElink 1000 eye-tracker system (SR Research Ltd.) with 500 ms sampling rate. Participants were instructed not to blink during the presentation of the options. Pre-processing of the pupil data was performed in MATLAB (version 2017a, MathWorks, Natick, USA). Data indicating eye blinks were replaced using linear interpolation. The data were visually inspected to ensure that all artefacts had been successfully removed. Pupil data were z-transformed within each run to control for variability across runs and across subjects. Pupil rate of dilation (unit: z/s), our measure of arousal, was calculated by subtracting pupil size at button response from pupil size at stimulus onset, divided by RT. Pre-trial pupil baseline level (PBL) was calculated by averaging pupil size from 500ms - 1ms before stimulus onset.

To ensure constant screen luminance level, we kept roughly the same number of pixels throughout the events by replacing the text indicating reward and effort levels with a series of Xs and by using text hues that were isoluminant to the grey background (RGB grey: 178.5, 178.5, 178.5; green: 50, 100, 10; purple: 118, 60, 206; blue: 53 77 229). Ensuring readability, we selected these hues out of 17 theoretically isoluminant hues where relative luminance was calculated as a linear combination of the red, green, and blue components based on the formula: Y = 0.2126 R + 0.7152 G + 0.0722 B. This formula follows the function that green light contributes the most to perceived intensity while blue contributes the least (Stokes, et al.; https://www.w3.org/Graphics/Color/sRGB). Green was always fixed as the base hue and blue and purple were randomly assigned trial-by-trial to highlight the selected offer (Fig. 1A).

Additionally, in a control experiment, we recorded luminance-driven pupil dilation without any cognitive task during presentation of fixation screens with a series of Xs as fixation period and Ys to replace the text that would have indicated the effort and reward levels in the main experiment, each period lasting for 3 s. Participants were instructed to keep their eyes open but were not required to press any key. Just like in the main experiment, green was the base hue during fixation whereas blue and purple were used to highlight the text on one side of the screen. All stimuli were in the same text format as in the main task (Fig. 1). Order of hue and side assignment were all counterbalanced and pseudorandomised. We found no difference in mean pupil diameter during the presentation of these control stimuli in different hues, confirming that the pupil response in the main task was not driven by differences in text luminance (Fig. S1).

### fMRI acquisition and analysis

Functional imaging was performed on a Philips Achieva 3T whole-body MR scanner equipped with a 32-channel MR head coil. Each experimental run contained 225-244 volumes (voxel size, 3×3×3 mm_3_; 0.5 mm gap; matrix size, 80×78 (FoV: [240 140 (FH) 240]; TR/TE 2334/30 ms; flip angle, 90°; parallel imaging factor, 1.5; 40 slices acquired in ascending order for full coverage of the brain). We also acquired T1-weighted multislice gradient-echo B0 scans which were used for correction of deformations (voxel size, 3 × 3 ×3 mm^3^; 0.75 mm gap; matrix size, 80×80; TR/TE1/TE2 ⫽ 400/4.3/7.4 ms; flip angle, 44°; parallel imaging; 40 slices). Additionally, we acquired a high-resolution T1-weighted 3D fast-field echo structural scan used for image registration during postprocessing (170 sagittal slices; matrix size, 256×256; voxel size, 1×1×1 mm^3^; TR/TE/TI ⫽ 8.3/3.9/1098 ms).

We used Statistical Parametric Mapping (SPM12; Wellcome Trust Centre for Neuroimaging, London, http://www.fil.ion.ucl.ac.uk/spm) for imaging analyses. Five preprocessing steps included (1) realignment and unwarping, (2) slice-timing correction, (3) coregistration and normalization, (4) smoothing, and (5) correction for physiological noise. First, we re-aligned all functional volumes to the first volume to correct for inter-scan movement. Images were unwarped using field maps to remove unwanted variance due to field inhomogeneity (Andersson et al., 2001). Second, unwarped functional images were slice-time corrected (to the acquisition time of the middle slice). Third, each subjects’ T1 image was co-registered (as reference image) with the mean functional image (as source image) using segmentation parameters performed on both images (Ashburner and Friston, 2004). These images were then normalized using the inverse deformation procedure and spatially re-sampled to 3 mm isotropic voxels. Fourth, all images were smoothed using a Gaussian kernel (FWHM 8mm). Finally, we performed correction for physiological noise via RETROICOR (Glover et al., 2000; Hutton et al., 2011) using Fourier expansions of different order for the estimated phases of cardiac pulsation (3rd order), respiration (4th order) and cardio‐respiratory interactions (1st order) (Hutton et al., 2011). We created the corresponding confound regressors using the PhysIO Toolbox (Kasper et al., 2009) (https://www.translationalneuromodeling.org/tapas).

We performed random-effect, event-related statistical analyses. For each subject, we first computed a statistical general linear model (GLM) by convolving series of stick functions (time-locked to the stimulus onsets and with the trial-wise RT as each event’s duration) with the canonical hemodynamic response functions and their first derivatives (temporal derivative). We also added to these GLMs 18 physiological regressors and 6 motion parameters. At the second level, we then tested the significance of subject-specific effects (as tested by t-contrasts at the first level) across the population. For these analyses, we used a grey matter mask as an explicit mask, created by averaging across subjects and smoothing (8mm) all participants’ normalized grey matter images (wc1*.nii) from the ‘segment’ procedure.

We built two first level GLMs without any orthogonalization. To identify unique variance associated with each of our trial parameters, we generated GLM1 using the stimulus onset as a single regressor with choice (1: effortful, −1: non-effortful), reward and effort levels of the effortful option, RT, pupil rate, and effort-by-choice (all non-binary variables were z-scored) as trial-wise parametric modulators. We then entered the contrast images of each parametric modulator vs baseline into second level one-sample t-tests. To illustrate the effort-by-choice interaction effect, we generated GLM2 with two regressors containing the stimulus onsets for choose effortful and choose non-effortful trials. Each regressor contained reward and effort levels of the effortful option, RT and pupil rate (all z-scored) as trial-wise parametric modulators. We then entered the contrast images of the effort parametric modulator for [choose effortful > choose non-effortful] into second level one-sample t-tests.

### Statistics

Statistical analyses for behavioral and pupil data were done with MATLAB 2017 (www.mathworks.com). We conducted (multiple) logistic or linear regressions separately for each participant and entered the regression weights of each predictor from all participants into a one-sample t-test. All continuous predictors were z-scored across trials within each participant. This approach allows for the intercept (constant) to vary across participants. Goodness-of-fit is the adjusted R_2_ for regressions. We used robust regression to evaluate the association between two variables. All statistical tests were two-tailed. For inference about the brain data, we used a cluster-defining threshold of *p*<0.001 and only report suprathreshold voxels that survive cluster-level family-wise error (FWE) corrected *p*<0.05.

## Acknowledgments

**General:** The authors thank Yoojin Lee and Zoltan Nagy for assistance in MRI optimisation, Karl Treiber and Miguel A. Garcia for assistance in data collection, and our participants for their voluntary participation.

## Author contributions

I.K. and C.C.R. conceived and designed the experiment. I.K. carried out the experiment. I.K. conducted all analyses with input from M.G. and C.C.R.. M.G. provided analytical software. I.K., M.G., and C.C.R interpreted the results and wrote the manuscript.

## Data availability

All raw and processed data, as well as the code to reproduce all analyses and figures will be made available on github or the OSF upon publication.

## Supplementary Materials and Methods

### Controlling for luminance-driven pupil response

To rule out brightness-induced pupil dilation and to validate our selection of theoretically isoluminant stimuli, we recorded pupil response during a control experiment at the end of the fMRI scan. Here, the same participants received similar visual stimulation as in the main experiment, but without informative cues or any need for making a choice. Participants were firs t presented with the same fixation screen (Fig 2A; screen with “XXX”) with letters written in green for 3 s. This was followed by the same screen but with all Xs replaced by Ys, and in either one of the sides (counterbalanced), the letters were printed in either purple or blue ink (to mimic the visual change found in the main experiment) for another 3 s period. All three color hues are theoretically isoluminant, as described in the Methods section. There were 20 trials for each side and each hue, summing to 80 trials. We confirm that indeed the hue selection in a task without any reward-effort decision making did not evoke meaningful luminance-driven pupil variance (Fig S1). First, the scale of pupil response variance in the main task was at least 6 times larger than that in the control experiment. Second, if any, the deflection in pupil response to cue onset was negative, as opposed to that found in the main task. Third, this control experiment revealed no difference in averaged pupil size across the entire stimulus duration between the two isoluminant hues (purple and blue) used in the main task, paired-samples t-test: *t*(46)=0.29, *p*=0.76 (2 missing data). These results confirm that the pupil dilation observed in the main task was primarily driven by meaningful cognitive considerations provoked by the choice task, in this case by effort-reward tradeoffs, and not by task-irrelevant physical differences in the stimuli.

**Fig. S1.**
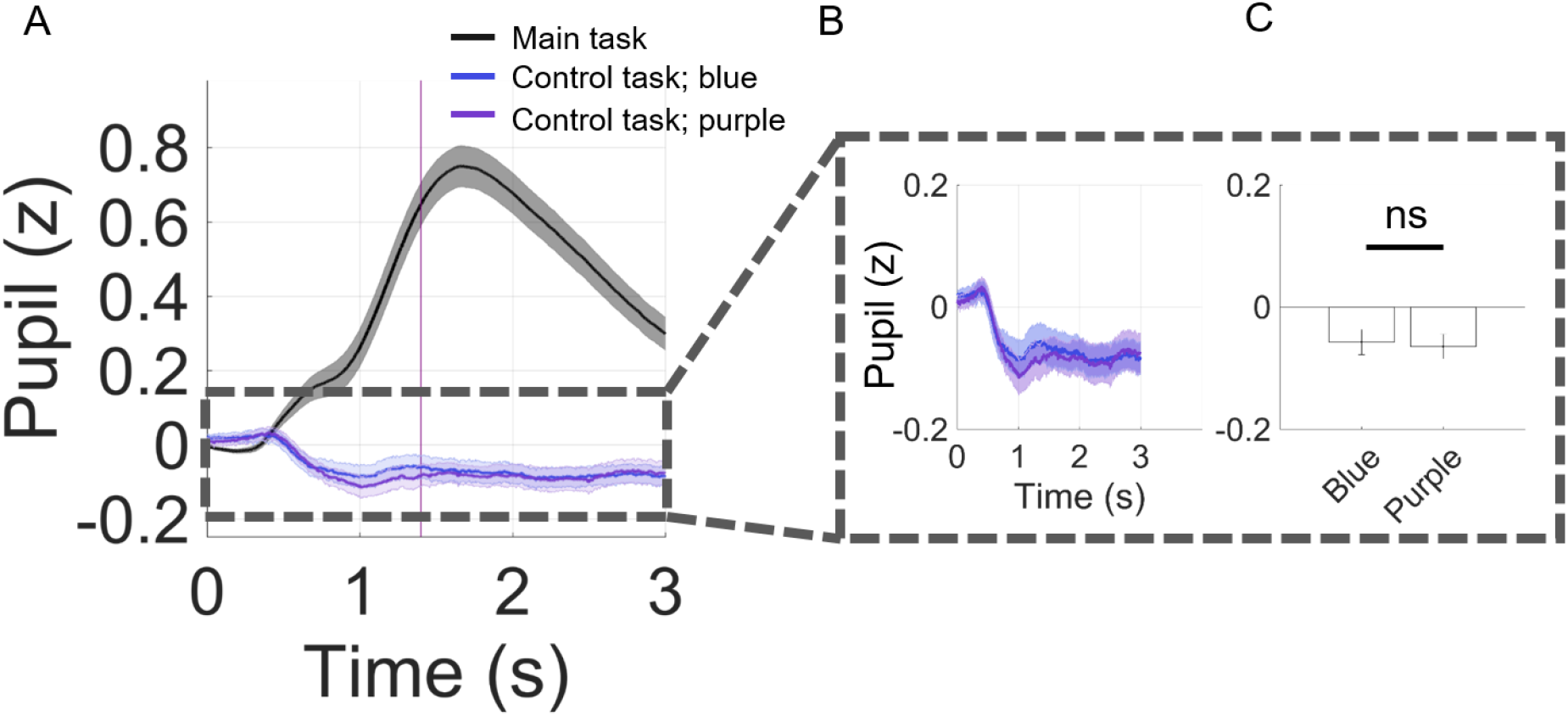
Pupil during main versus control experiment. A) Pupil time course in main and in control task for stimuli in blue and purple ink, subtracted by pupil baseline level (PBL). Inset: B) Zoomed-in pupil time course and C) averaged pupil size across 3 s, showing no difference in pupil responses between blue and purple. Bar plots display means ± 1 standard error of the mean (SEM).

### Controlling for other variables in extended regression of choice (Fig 2D)

Here we supplement the statistical results of the regression of choice reported in the main text (Fig 2D). In this extended regression, we also accounted for variables including RT (*t*_RT_(48)=-3.40, *p*=0.0013), pupil baseline level (PBL), *t*_PBL_(48)=0.25, *p*=0.80, and many others (*t*_pupil_rate_(48)=-1.02, *p*=0.31; *t*_reward*PBL_(48)=0.22, *p*=0.82; *t*_ef f ort*PBL_(48)=-0.31, *p*=0.75; *t*_reward*ef f ort*PBL_(48)=-0.61, *p*=0.54; *t*_reward*pupil_rate_(48)=-0.78, *p*=0.44; *t*_reward*ef f ort*pupil_rate_(48)=1.21, *p*=0.23; *t*_constant_(48)=4.37, *p*=0.0001). Importantly, the extended regression had a higher model-fit (adjusted *R*-squared) than the standard regression that only contained reward, effort, and reward-by-effort, *t*(48)=5.35, *p*<0.0001, suggesting that pupil measures together with other task parameters such as reward, effort, and response time, can explain choice above and beyond the ‘standard’ option attributes (reward and effort).

### Calculation of effort slope

To calculate the effort slope in all our analyses (e.g., Fig 3), for each subject we first averaged the pupil rate (z-scored within subject) in each of the 6 effort levels separately for trials where subjects chose the effortful option and those where they chose the non-effortful option. We then ran a simple regression of the averaged pupil rate on effort levels (levels 4-9), separately for each choice outcome. Without any missing data, the effort slope should be estimated based on 6 pairs of data points. However, choice was clearly affected by effort level (Fig 1C), thus one concern is that for some subjects, there might have been too many empty cells (e.g., if options with effort levels 7-9 were never selected by a participant). If this were the case then there would have unequal number of data points to estimate the effort slope in one choice outcome versus another. To address this concern, we found that on average there were > 5 pairs of data points in both choice outcomes (*M*_non-ef f ortful_=5.59, SD=0.67, *M*_ef fortful_=5.61 SD=0.7), and importantly there was no significant difference between the two choice outcomes, *t*(48)=0.13, *p*=0.89. This result assured us that the estimation of effort slopes between the two choice outcomes was comparable.

### Control analysis for pupil baseline level (PBL)

To investigate how other aspects of the arousal system function may relate to choice in our experimental design, we conducted a whole set of other analyses. First, we examined choice proportions as a function of PBL median/tertile/quartile splits. We did not find any choice differences across PBL bins, *F*s<1.2, *p*s>0.3 (Fig S2A). Second, we ran a logistic regression of choice on PBL, RT, reward, effort, and the interactions. We found no effect of PBL or any interactions with PBL (Fig S2B). Third, we inspected whether regressing out influences of previous trial from PBL would improve regression of choice of the current trial. To do this, we first ran a linear regression of the current trial’s PBL with reward, effort, choice, RT, and ITI of previous trial (t-1) as regressors. Then we took the residual variance of this regression and used it as a regressor together with RT, reward, effort, reward-by-effort interaction to fit choice of current trial. This analysis shows no significant effect of the residual PBL (PBL*) on explaining choice of current trial (Fig S2C). Together, these analyses show no contribution of background (tonic) arousal states to choice rate, suggesting that the results reported in the main text were specific to effort-specific representations during the decision process (within-trial).

**Fig. S2.**
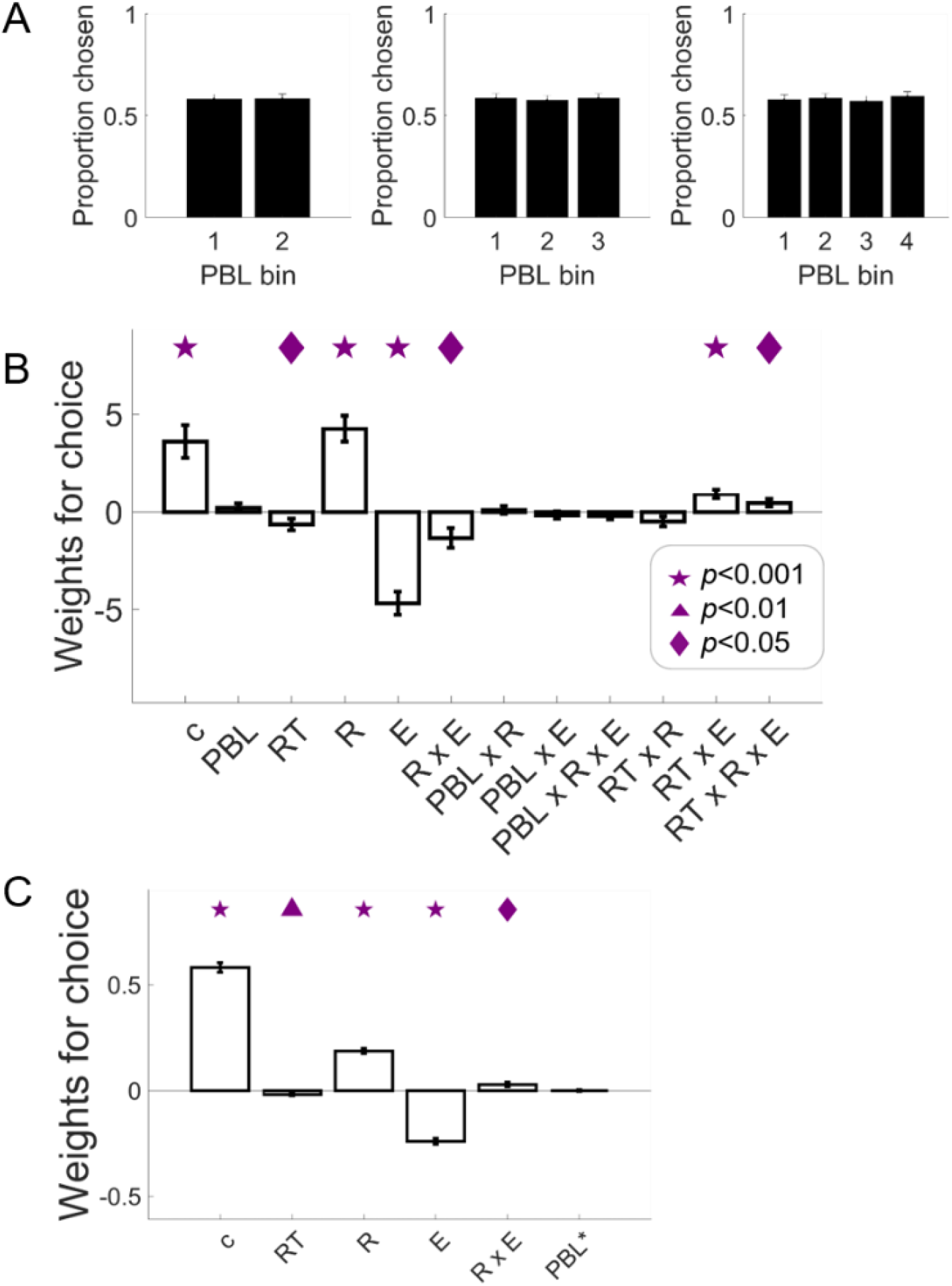
No effects of endogenous arousal fluctuations on choice rate. A) Choice proportion for the effortful option as a function of pre-trial pupil baseline level (PBL) bins. Bar plots display means ± 1 standard error of the mean (SEM). B) Weights of logistic regression of choice on reward, effort, RT, PBL, and the interactions. B Weights of logistic regression of choice on reward, effort, RT, and residual variance of PBL after regressing out influences from previous trial (t-1). Bar plots display means ± 1 standard error of the mean (SEM). Abbreviations: c=constant, PBL=pupil baseline levels, RT=reaction time, R=reward levels, E=effort levels, PBL*=residual PBL.

In addition, we also directly tested for an effort-by-choice effect in PBL (Fig S3), revealing a non-significant choice difference (effortful vs non-effortful) of the effort slopes in PBL, *t*(48)=0.45, *p*=0.65. The behavioral measure of effort sensitivity was not significantly associated with the effort-by-choice effect either, *robust regression b*(47)=0.35, *p*=0.39. These results confirm that the choice-modulated effort representations reported in the main text are primarily expressed in how fast the pupil dilates but not in endogenous pre-trial pupil fluctuations.

**Fig. S3.**
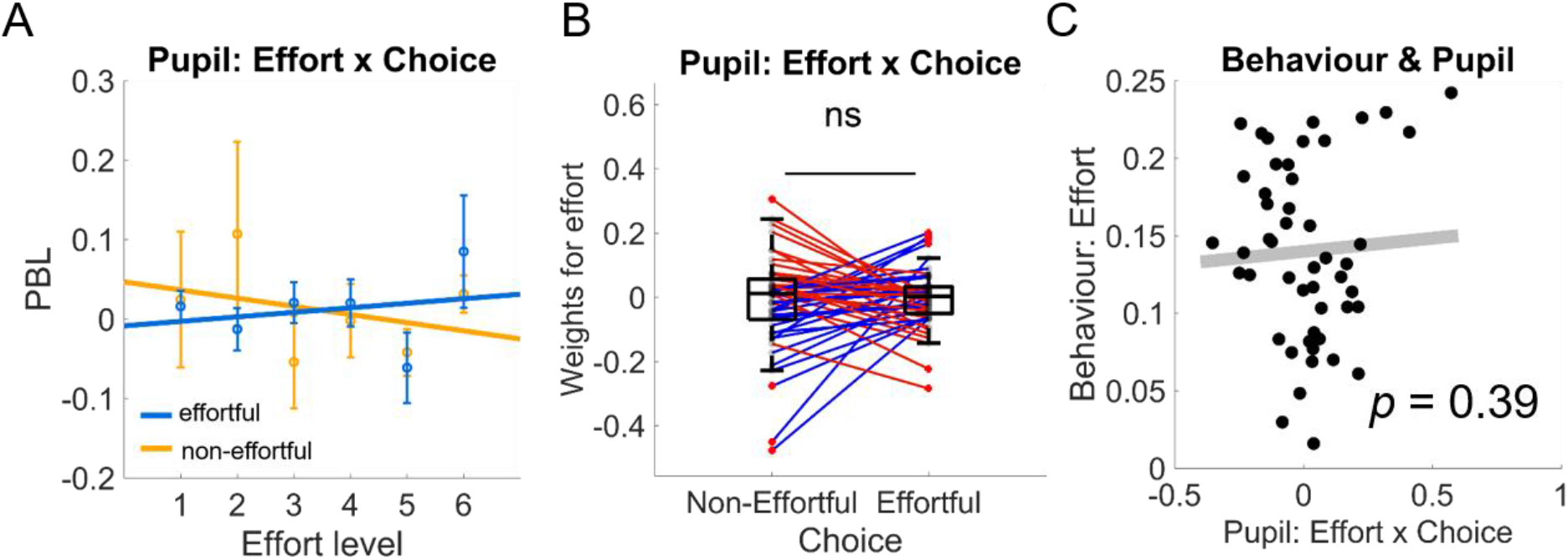
No evidence for energization signals in pupil baseline level (PBL). Non-significant effort-by-choice interaction results and non-significant correlation with behavioral effort sensitivity for PBL. Bar plots display means ± 1 standard error of the mean (SEM).

### Control analysis for effects of value (Fig 3)

To rule out the alternative explanation that pupil rate in this experiment could be simply signalling value, we tested for a reward-by-choice effect in pupil rate (Fig S5), revealing a non-significant choice difference (effortful vs non-effortful) of the reward slopes, *t*(48)=0.22, *p*=0.82. The behavioral measure of reward sensitivity was not significantly associated with the reward-by-choice effect either, *robust regression b*(47)=0.54, *p*=0.22. Together with the analyses on pupil rate residuals reported in the main text (Figs S4), these results confirm the pupil rate’s role in anticipated energization, signaling effort amounts that one has committed to rather than signaling reward value.

**Fig. S4.**
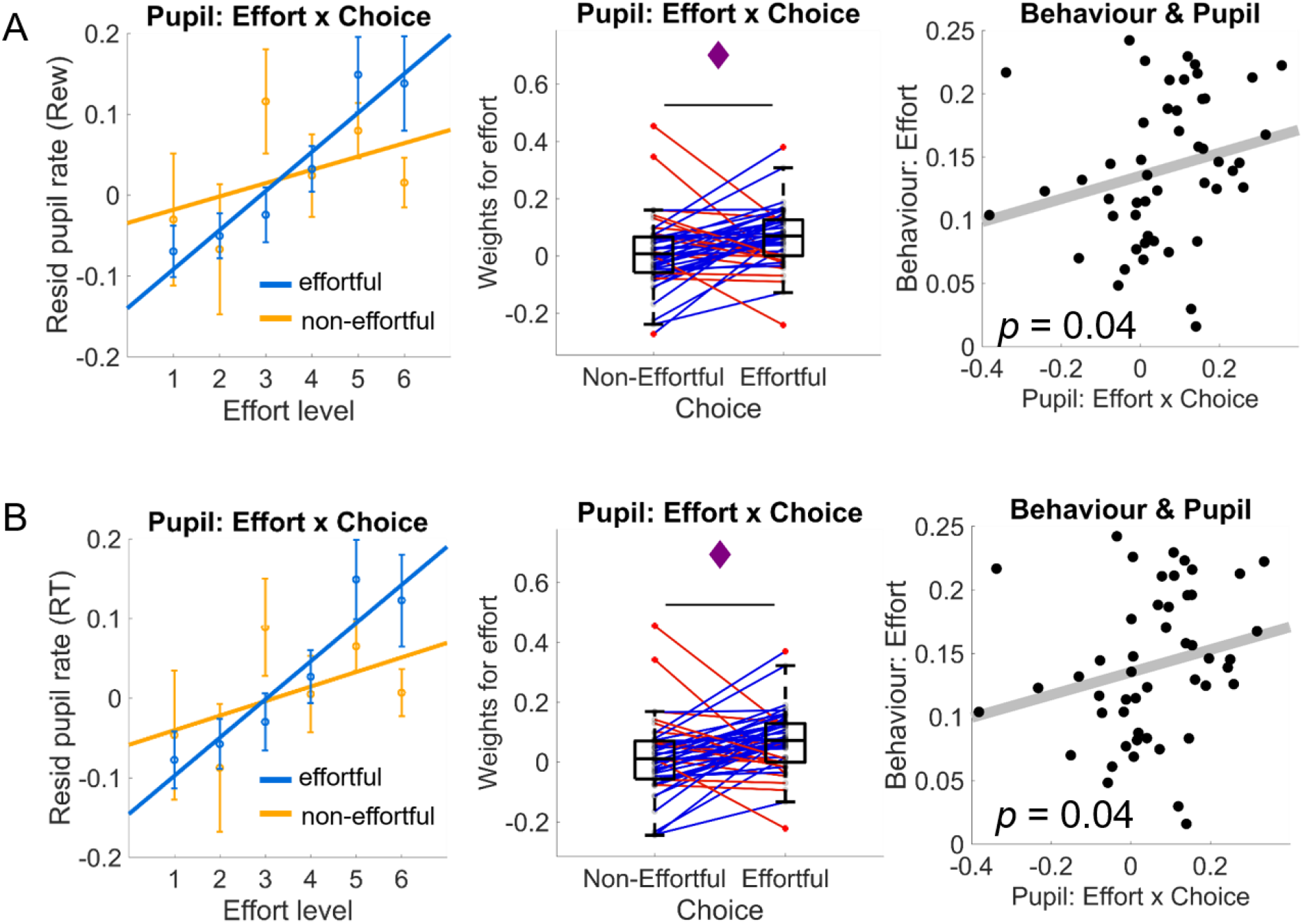
Energization signal in pupil rate is independent of reward value and RT. Analyses with residual pupil rate after regressing out the effect of reward (A) and RT (B). We replicated the effects reported in Fig 3, showing significant effort-by-choice interaction results and correlation with behavioral effort sensitivity for residual pupil rate after regressing out (one at a time) the effect of reward and RT. Bar plots display means ± 1 standard error of the mean (SEM). See main text for statistical results.

**Fig. S5.**
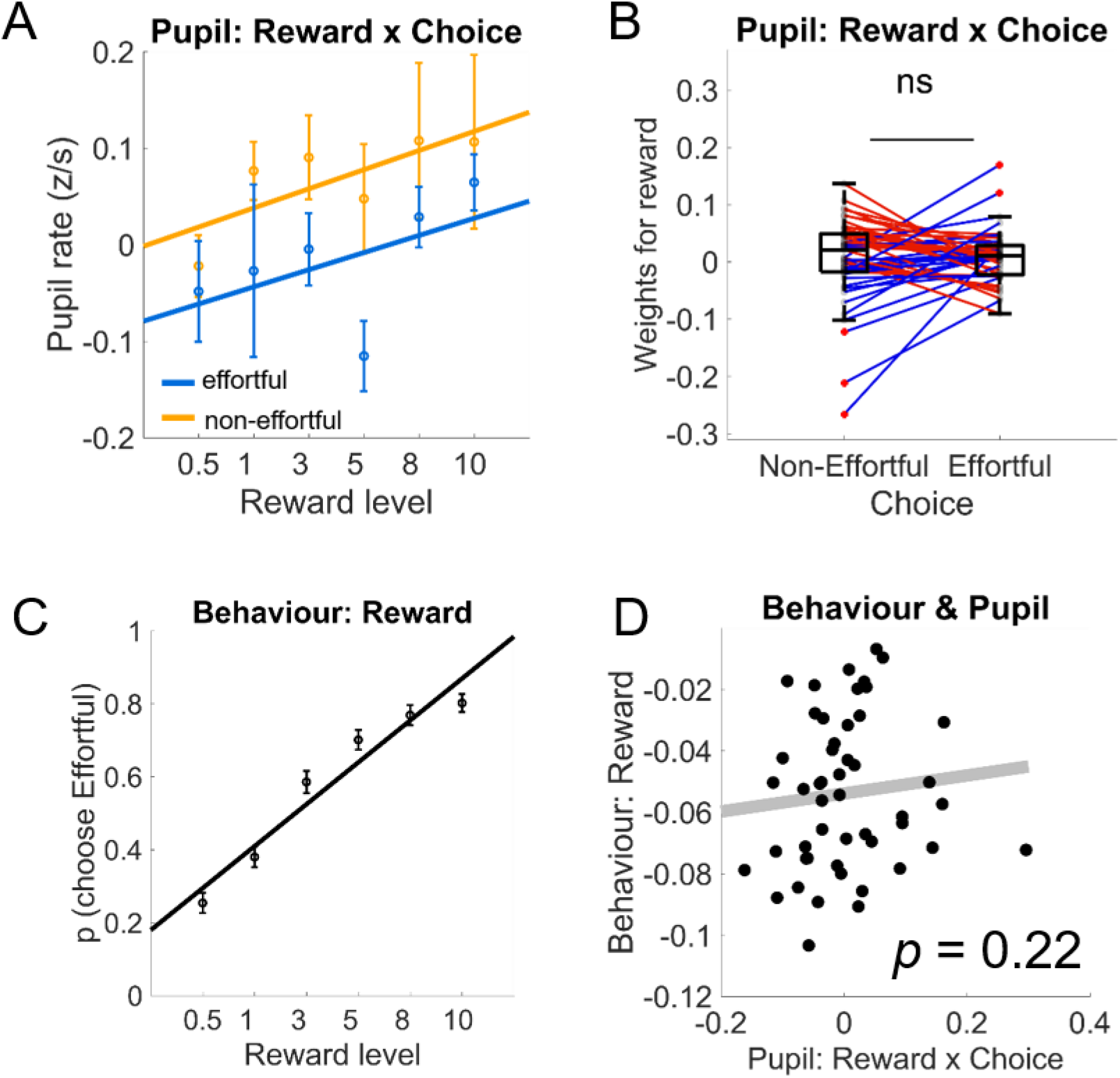
No evidence for choice-modulated reward signals in pupil rate. Non-significant **r**eward-by-choice interaction results and non-significant correlation with behavioral reward sensitivity in pupil rate. Bar plots display means ± 1 standard error of the mean (SEM).

## References

Aridan, Nadav, Nicholas J. Malecek, Russell A. Poldrack, and Tom Schonberg. 2019. “Neural Correlates of Effort-Based Valuation with Prospective Choices.” NeuroImage 185 (July 2018): 446–54. https://doi.org/10.1016/j.neuroimage.2018.10.051.

Arnsten, A. F.T., and P. S. Goldman-Rakic. 1984. “Selective Prefrontal Cortical Projections to the Region of the Locus Coeruleus and Raphe Nuclei in the Rhesus Monkey.” Brain Research 306 (1–2): 9–18. https://doi.org/10.1016/0006-8993(84)90351-2.

Arulpragasam, Amanda R., Jessica A. Cooper, Makiah R. Nuutinen, and Michael T. Treadway. 2018. “Corticoinsular Circuits Encode Subjective Value Expectation and Violation for Effortful Goal-Directed Behavior.” Proceedings of the National Academy of Sciences 115 (22): E5233–42. https://doi.org/10.1073/pnas.1800444115.

Aston-Jones, Gary, and Jonathan D. Cohen. 2005. “An Integrative Theory of Locus Coeruleus - Norepinephrine Function: Adaptive Gain and Optimal Performance.” Annual Review of Neuroscience 28 (1): 403–50. https://doi.org/10.1146/annurev.neuro.28.061604.135709.

Bartra, Oscar, Joseph T. McGuire, and Joseph W Kable. 2013. “The Valuation System: A Coordinate-Based Meta-Analysis of BOLD FMRI Experiments Examining Neural Correlates of Subjective Value.” NeuroImage 76 (August): 412–27. https://doi.org/10.1016/j.neuroimage.2013.02.063.

Bautista, Luis M, Joost Tinbergen, and Alex Kacelnik. 2001. “To Walk or to Fly? How Birds Choose among Foraging Modes.” Proc. Natl. Acad. Sci. U.SA. 98: 1089–94.

Beierholm, Ulrik, Marc Guitart-Masip, Marcos Economides, Rumana Chowdhury, Emrah Düzel, Ray Dolan, and Peter Dayan. 2013. “Dopamine Modulates Reward-Related Vigor.” Neuropsychopharmacology 38 (8): 1495–1503. https://doi.org/10.1038/npp.2013.48.

Benoit, Roland G, Sam J Gilbert, and Paul W Burgess. 2011. “A Neural Mechanism Mediating the Impact of Episodic Prospection on Farsighted Decisions.” Journal of Neuroscience 31 (18): 6771–79. https://doi.org/10.1523/JNEUROSCI.6559-10.2011.

Bernacer, Javier, Ivan Martinez-Valbuena, Martin Martinez, Nuria Pujol, Elkin Luis, David Ramirez-Castillo, and Maria A. Pastor. 2019. “Neural Correlates of Effort-Based Behavioral Inconsistency.” Cortex 113 (Imim): 96–110. https://doi.org/10.1016/j.cortex.2018.12.005.

Berwian, IM;, J; Wenzel, AGE; Collins, E; Seifritz, KE; Stephan, H; Walter, and QJM; Huys. 2020. “Computational Mechanisms of Effort and Reward Decisions in Depression and Their Relationship to Relapse after Antidepressant Discontinuation.” JAMA Psychiatry, In Press. https://doi.org/10.1001/jamapsychiatry.2019.4971.

Borderies, Nicolas, Pauline Bornert, Sophie Gilardeau, and Sebastien Bouret. 2020. “Pharmacological Evidence for the Implication of Noradrenaline in Effort.” PLoS Biology 18 (10): 1–26. https://doi.org/10.1101/714923.

Bulley, Adam, and Matthew J. Gullo. 2017. “The Influence of Episodic Foresight on Delay Discounting and Demand for Alcohol.” Addictive Behaviors. https://doi.org/10.1016/j.addbeh.2016.11.003.

Burke, Christopher J, Christian Brünger, Thorsten Kahnt, Soyoung Q Park, and Philippe N Tobler. 2013. “Neural Integration of Risk and Effort Costs by the Frontal Pole: Only upon Request.” Journal of Neuroscience 33 (4): 1706–13. https://doi.org/10.1523/JNEUROSCI.3662-12.2013.

Chandler, Daniel J., Carolyn S. Lamperski, and Barry D. Waterhouse. 2013. “Identification and Distribution of Projections from Monoaminergic and Cholinergic Nuclei to Functionally Differentiated Subregions of Prefrontal Cortex.” Brain Research 1522 (July): 38–58. https://doi.org/10.1016/j.brainres.2013.04.057.

Chong, Trevor T.-J., Matthew Apps, Kathrin Giehl, Annie Sillence, Laura L. Grima, and Masud Husain. 2017. “Neurocomputational Mechanisms Underlying Subjective Valuation of Effort Costs.” Edited by Ben Seymour. PLOS Biology 15(2): e1002598. https://doi.org/10.1371/journal.pbio.1002598.

Cole, Michael W., Jeremy R. Reynolds, Jonathan D. Power, Grega Repovs, Alan Anticevic, and Todd S. Braver. 2013. “Multi-Task Connectivity Reveals Flexible Hubs for Adaptive Task Control.” Nature Neuroscience 16 (9): 1348–55. https://doi.org/10.1038/nn.3470.

Dalsass, M., Sanford Kiser, Margaret Mèndershausen, and D. C. German. 1981. “Medial Prefrontal Cortical Projections to the Region of the Dorsal Periventricular Catecholamine System.” Neuroscience. https://doi.org/10.1016/0306-4522(81)90149-4.

Dassen, Fania C.M. M, Anita Jansen, Chantal Nederkoorn, and Katrijn Houben. 2016. “Focus on the Future: Episodic Future Thinking Reduces Discount Rate and Snacking.” Appetite 96: 327–32. https://doi.org/10.1016/j.appet.2015.09.032.

Dosenbach, Nico U.F., Damien A. Fair, Alexander L. Cohen, Bradley L. Schlaggar, and Steven E. Petersen. 2008. “A Dual-Networks Architecture of Top-down Control.” Trends in Cognitive Sciences 12 (3): 99–105. https://doi.org/10.1016/j.tics.2008.01.001.

Ebitz, R. Becket, and Michael L. Platt. 2015. “Neuronal Activity in Primate Dorsal Anterior Cingulate Cortex Signals Task Conflict and Predicts Adjustments in Pupil-Linked Arousal.” Neuron 85 (3): 628–40. https://doi.org/10.1016/j.neuron.2014.12.053.

Fleming, Stephen Michael, and Nathaniel D Daw. 2017. “Self-Evaluation of Decision-Making: A General Bayesian Framework for Metacognitive Computation.” Psychological Review 124 (1): 91–114. https://doi.org/10.1037/rev0000045.

Gaesser, Brendan, Kerri Keeler, and Liane Young. 2018. “Moral Imagination: Facilitating Prosocial Decision-Making through Scene Imagery and Theory of Mind.” Cognition 171 (February): 180–93. https://doi.org/10.1016/j.cognition.2017.11.004.

Gee, Jan Willem de, Olympia Colizoli, Niels A. Kloosterman, Tomas Knapen, Sander Nieuwenhuis, and Tobias H. Donner. 2017. “Dynamic Modulation of Decision Biases by Brainstem Arousal Systems.” ELife 6 (Lc): 1–36. https://doi.org/10.7554/eLife.23232.

Gee, Jan Willem de, Tomas Knapen, and Tobias H Donner. 2014. “Decision-Related Pupil Dilation Reflects Upcoming Choice and Individual Bias.” Proceedings of the National Academy of Sciences of the United States of America 111 (5): E618–25. https://doi.org/10.1073/pnas.1317557111.

Gelbard-Sagiv, Hagar, Efrat Magidov, Haggai Sharon, Talma Hendler, and Yuval Nir. 2018. “Noradrenaline Modulates Visual Perception and Late Visually Evoked Activity.” Current Biology 28 (14): 2239–2249.e6. https://doi.org/10.1016/j.cub.2018.05.051.

Glimcher, Paul W. 2009. “Choice: Towards a Standard Back-Pocket Model.” In Neuroeconomics: Decision Making and the Brain, edited by Paul W. Glimcher, Colin F. Camerer, Ernst Fehr, and Russell A. Poldrack, First, 503–21. London: Academic Press. https://doi.org/10.1016/B978-0-12-374176-9.00032-4.

Goodale, Melvyn A., and A. David Milner. 2018. “Separate Visual Pathways for Perception and Action.” Human Perception: Institutional Performance and Reform in Australia, no. I: 123–28. https://doi.org/10.4324/9781351156288-16.

Grueschow, Marcus, Birgit Kleim, and Christian C. Ruff. 2020. “Role of the Locus Coeruleus Arousal System in Cognitive Control.” Journal of Neuroendocrinology, no. July: 1–11. https://doi.org/10.1111/jne.12890.

Hassabis, Demis, and Eleanor A. Maguire. 2007. “Deconstructing Episodic Memory with Construction.” Trends in Cognitive Sciences. https://doi.org/10.1016/j.tics.2007.05.001.

Hauser, Tobias U., Eran Eldar, and Raymond J. Dolan. 2017. “Separate Mesocortical and Mesolimbic Pathways Encode Effort and Reward Learning Signals.” Proceedings of the National Academy of Sciences, 201705643. https://doi.org/10.1073/pnas.1705643114.

Heron, C. Le, M. A.J. Apps., and M. Husain. 2018. “The Anatomy of Apathy: A Neurocognitive Framework for Amotivated Behaviour.” Neuropsychologia 118 (May 2017): 54–67. https://doi.org/10.1016/j.neuropsychologia.2017.07.003.

Hockey, G. Robert, J. 1997. “Compensatory Control in the Regulation of Human Performance under Stress and High Workload: A Cognitive-Energetical Framework.” Biological Psychology 45 (1–3): 73–93. https://doi.org/10.1016/S0301-0511(96)05223-4.

Hull, C. L. 1943. Principles of Behavior: An Introduction to Behavior Theory. New York: Appleton-Century-Crofts.

Husain, Masud, and Jonathan P. Roiser. 2018. “Neuroscience of Apathy and Anhedonia: A Transdiagnostic Approach.” Nature Reviews Neuroscience 19 (8): 470–84. https://doi.org/10.1038/s41583-018-0029-9.

Jahn, Caroline I., Sophie Gilardeau, Chiara Varazzani, Bastien Blain, Jerome Sallet, Mark E. Walton, and Sebastien Bouret. 2018. “Dual Contributions of Noradrenaline to Behavioural Flexibility and Motivation.” Psychopharmacology 235 (9): 2687–2702. https://doi.org/10.1007/s00213-018-4963-z.

Joshi, Siddhartha, Yin Li, Rishi M. Kalwani, and Joshua I. Gold. 2016. “Relationships between Pupil Diameter and Neuronal Activity in the Locus Coeruleus, Colliculi, and Cingulate Cortex.” Neuron 89 (1): 221–34. https://doi.org/10.1016/j.neuron.2015.11.028.

Kiani, Roozbeh, Leah Corthell, and Michael N. Shadlen. 2014. “Choice Certainty Is Informed by Both Evidence and Decision Time.” Neuron. https://doi.org/10.1016/j.neuron.2014.12.015.

Klein-Flügge, Miriam Cornelia, Steven W. Kennerley, Karl Friston, and Sven Bestmann. 2016. “Neural Signatures of Value Comparison in Human Cingulate Cortex during Decisions Requiring an Effort-Reward Trade-Off.” Journal of Neuroscience 36 (39): 10002–15. https://doi.org/10.1523/JNEUROSCI.0292-16.2016.

Kurniawan, Irma Triasih, Marc Guitart-Masip, P. Dayan, and Raymond J. Dolan. 2013. “Effort and Valuation in the Brain: The Effects of Anticipation and Execution.” Journal of Neuroscience 33 (14): 6160–69. https://doi.org/10.1523/JNEUROSCI.4777-12.2013.

Kurniawan, Irma Triasih, Marc Guitart-Masip, and Raymond J. Dolan. 2011. “Dopamine and Effort-Based Decision Making.” Frontiers in Decision Neuroscience 5 (81): 1–10. https://doi.org/10.3389/fnins.2011.00081.

Kurniawan, Irma Triasih, Ben Seymour, Deborah Talmi, Wako Yoshida, Nick Chater, and Raymond J. Dolan. 2010. “Choosing to Make an Effort: The Role of Striatum in Signaling Physical Effort of a Chosen Action.” Journal of Neurophysiology 104 (1): 313–21. https://doi.org/10.1152/jn.00027.2010.

McGinley, Matthew J., Martin Vinck, Jacob Reimer, Renata Batista-Brito, Edward Zagha, Cathryn R. Cadwell, Andreas S. Tolias, Jessica A. Cardin, and David A. McCormick. 2015. “Waking State: Rapid Variations Modulate Neural and Behavioral Responses.” Neuron 87 (6): 1143–61. https://doi.org/10.1016/j.neuron.2015.09.012.

McGuire, Joseph T., and Matthew M Botvinick. 2010. “Prefrontal Cortex, Cognitive Control, and the Registration of Decision Costs.” Proceedings of the National Academy of Sciences of the United States of America 107 (17): 7922–26. https://doi.org/10.1073/pnas.0910662107.

Meyniel, Florent, Guy M. Goodwin, J. F. William Deakin, Corinna Klinge, Christine Macfadyen, Holly Milligan, Emma Mullings, Mathias Pessiglione, and Raphaël Gaillard. 2016. “A Specific Role for Serotonin in Overcoming Effort Cost.” ELife 5 (NOVEMBER2016): 1–18. https://doi.org/10.7554/eLife.17282.

Meyniel, Florent, Claire Sergent, Lionel Rigoux, Jean Daunizeau, and Mathias Pessiglione. 2013. “Neurocomputational Account of How the Human Brain Decides When to Have a Break.” Proceedings of the National Academy of Sciences of the United States of America 110 (7): 2641–46. https://doi.org/10.1073/pnas.1211925110.

Moret, Chantal, and Mike Briley. 2011. “The Importance of Norepinephrine in Depression.” Neuropsychiatric Disease and Treatment 7 (SUPPL.): 9–13. https://doi.org/10.2147/NDT.S19619.

Murphy, Peter R., Joachim Vandekerckhove, and Sander Nieuwenhuis. 2014. “Pupil-Linked Arousal Determines Variability in Perceptual Decision Making.” PLoS Computational Biology 10 (9). https://doi.org/10.1371/journal.pcbi.1003854.

Niv, Yael, Nathaniel D Daw, and Peter Dayan. 2005. “How Fast to Work : Response Vigor, Motivation and Tonic Dopamine.” In Neural Information Processing Systems, edited by Y Weiss, B Scholkopf, and J Platt, 1019–26. MIT Press.

Nutt, David, Koen Demyttenaere, Zoltan Janka, Trond Aarre, Michel Bourin, Pier Luigi Canonico, Jose Luis Carrasco, and Steven Stahl. 2007. “The Other Face of Depression, Reduced Positive Affect: The Role of Catecholamines in Causation and Cure.” Journal of Psychopharmacology 21 (5): 461–71. https://doi.org/10.1177/0269881106069938.

Ostlund, Sean B, K. M. Wassum, N. P. Murphy, Bernard W. Balleine, and N. T. Maidment. 2011. “Extracellular Dopamine Levels in Striatal Subregions Track Shifts in Motivation and Response Cost during Instrumental Conditioning.” Journal of Neuroscience 31 (1): 200–207. https://doi.org/10.1523/JNEUROSCI.4759-10.2011.

Paravlic, Armin H., Maamer Slimani, David Tod, Uros Marusic, Zoran Milanovic, and Rado Pisot. 2018. “Effects and Dose–Response Relationships of Motor Imagery Practice on Strength Development in Healthy Adult Populations: A Systematic Review and Meta-Analysis.” Sports Medicine 48 (5): 1165–87. https://doi.org/10.1007/s40279-018-0874-8.

Parvizi, Josef, Vinitha Rangarajan, William R Shirer, Nikita Desai, and Michael D Greicius. 2013. “Case Study The Will to Persevere Induced by Electrical Stimulation of the Human Cingulate Gyrus.” Neuron 80 (6): 1359–67. https://doi.org/10.1016/j.neuron.2013.10.057.

Pessiglione, Mathias, Fabien Vinckier, Sébastien Bouret, Jean Daunizeau, and Raphaël Le Bouc. 2018. “Why Not Try Harder? Computational Approach to Motivation Deficits in Neuro-Psychiatric Diseases.” Brain 141 (3): 629–50. https://doi.org/10.1093/brain/awx278.

Peters, Jan, and Christian Büchel. 2010. “Episodic Future Thinking Reduces Reward Delay Discounting through an Enhancement of Prefrontal-Mediotemporal Interactions.” Neuron 66 (1): 138–48. https://doi.org/10.1016/j.neuron.2010.03.026.

Pfaff, Donald W., Eugene M. Martin, and Donald Faber. 2012. “Origins of Arousal: Roles for Medullary Reticular Neurons.” Trends in Neurosciences 35 (8): 468–76. https://doi.org/10.1016/j.tins.2012.04.008.

Poe, Gina R, Stephen Foote, Oxana Eschenko, Joshua P Johansen, Sebastien Bouret, Gary Aston-Jones, Carolyn W Harley, et al. 2020. “Locus Coeruleus: A New Look at the Blue Spot.” Nature Reviews. Neuroscience. https://doi.org/10.1038/s41583-020-0360-9.

Porrino, Linda J., and Patricia S. Goldman‐Rakic. 1982. “Brainstem Innervation of Prefrontal and Anterior Cingulate Cortex in the Rhesus Monkey Revealed by Retrograde Transport of HRP.” Journal of Comparative Neurology 205 (1): 63–76. https://doi.org/10.1002/cne.902050107.

Prevost, Charlotte, Mathias Pessiglione, Elise Metereau, Marie-Laure Clery-Melin, and Jean-Claude Dreher. 2010. “Separate Valuation Subsystems for Delay and Effort Decision Costs.” Journal of Neuroscience 30 (42): 14080–90. https://doi.org/10.1523/JNEUROSCI.2752-10.2010.

Reimer, Jacob, Matthew J McGinley, Yang Liu, Charles Rodenkirch, Qi Wang, David A McCormick, and Andreas S Tolias. 2016. “Pupil Fluctuations Track Rapid Changes in Adrenergic and Cholinergic Activity in Cortex.” Nature Communications 7 (May): 13289. https://doi.org/10.1038/ncomms13289.

Salamone, John D., Mercè Correa, A. M. Farrar, and Susana Mingote. 2007. “Effort - Related Functions of Nucleus Accumbens Dopamine and Associated Forebrain Circuits.” Psychopharmacology 191: 461–82.

Saper, Clifford B., Patrick M. Fuller, Nigel P. Pedersen, Jun Lu, and Thomas E. Scammell. 2010. “Sleep State Switching.” Neuron 68 (6): 1023–42. https://doi.org/10.1016/j.neuron.2010.11.032.

Schmidt, Liane, Marie-Laure Cléry-Melin, Gilles Lafargue, Romain Valabrègue, Philippe Fossati, Bruno Dubois, and Mathias Pessiglione. 2009. “Get Aroused and Be Stronger: Emotional Facilitation of Physical Effort in the Human Brain.” Journal of Neuroscience 29 (30): 9450–57. https://doi.org/10.1523/JNEUROSCI.1951-09.2009.

Schneider, Max, Laura Leuchs, Michael Czisch, Philipp G. Sämann, and Victor I. Spoormaker. 2018. “Disentangling Reward Anticipation with Simultaneous Pupillometry / FMRI.” NeuroImage 178 (March): 11–22. https://doi.org/10.1016/j.neuroimage.2018.04.078.

Schultz, Wolfram. 2002. “Getting Formal with Dopamine and Reward.” Neuron 36 (2): 241–63. https://doi.org/10.1016/S0896-6273(02)00967-4.

Schultz, Wolfram, Peter Dayan, and P Read Montague. 1997. “A Neural Substrate of Prediction and Reward.” Science 275 (5306): 1593–99. http://www.ncbi.nlm.nih.gov/pubmed/9054347.

Schwarz, Lindsay A., Kazunari Miyamichi, Xiaojing J. Gao, Kevin T. Beier, Brandon Weissbourd, Katherine E. Deloach, Jing Ren, et al. 2015. “Viral-Genetic Tracing of the Input-Output Organization of a Central Noradrenaline Circuit.” Nature 524 (7563): 88–92. https://doi.org/10.1038/nature14600.

Shenhav, Amitai, Jonathan D Cohen, and Matthew M Botvinick. 2016. “Dorsal Anterior Cingulate Cortex and the Value of Control.” Nature Neuroscience 19 (10): 1286–91. https://doi.org/10.1038/nn.4384.

Skvortsova, V., S. Palminteri, and Mathias Pessiglione. 2014. “Learning To Minimize Efforts versus Maximizing Rewards: Computational Principles and Neural Correlates.” Journal of Neuroscience 34 (47): 15621–30. https://doi.org/10.1523/JNEUROSCI.1350-14.2014.

Steinmetz, Janina, Brittany M. Tausen, and Jane L. Risen. 2018. “Mental Simulation of Visceral States Affects Preferences and Behavior.” Personality and Social Psychology Bulletin 44 (3): 406–17. https://doi.org/10.1177/0146167217741315.

Takahashi, K., Y. Kayama, J. S. Lin, and K. Sakai. 2010. “Locus Coeruleus Neuronal Activity during the Sleep-Waking Cycle in Mice.” Neuroscience 169 (3): 1115–26. https://doi.org/10.1016/j.neuroscience.2010.06.009.

Tillage, Rachel P., Genevieve E. Wilson, L. Cameron Liles, Philip V. Holmes, and David Weinshenker. 2020. “Chronic Environmental or Genetic Elevation of Galanin in Noradrenergic Neurons Confers Stress Resilience in Mice.” The Journal of Neuroscience 40 (39): JN-RM-0973-20. https://doi.org/10.1523/jneurosci.0973-20.2020.

Varazzani, Chiara, a. San-Galli, S. Gilardeau, and S. Bouret. 2015. “Noradrenaline and Dopamine Neurons in the Reward/Effort Trade-Off: A Direct Electrophysiological Comparison in Behaving Monkeys.” Journal of Neuroscience 35 (20): 7866–77. https://doi.org/10.1523/JNEUROSCI.0454-15.2015.

Vinckier, Fabien, L. Rigoux, I.T. Kurniawan, C. Hu, Sacha Bourgeois-gironde, J. Daunizeau, and Mathias Pessiglione. 2018. “Sour Grapes and Sweet Victories : How Actions Shape Preferences.” Edited by Samuel J. Gershman. PLOS Computational Biology 15(1): e1006499. https://doi.org/10.1371/journal.pcbi.1006499.

Walton, Mark E., and Sebastien Bouret. 2019. “What Is the Relationship between Dopamine and Effort?” Trends in Neurosciences 42 (2): 79–91. https://doi.org/10.1016/j.tins.2018.10.001.

Wel, Pauline van der, and Henk van Steenbergen. 2018. “Pupil Dilation as an Index of Effort in Cognitive Control Tasks: A Review.” Psychonomic Bulletin & Review 25 (6): 2005–15. https://doi.org/10.3758/s13423-018-1432-y.

Xiang, Liyang, Antoine Harel, Hong Ying Gao, Anthony E. Pickering, Susan J. Sara, and Sidney I. Wiener. 2019. “Behavioral Correlates of Activity of Optogenetically Identified Locus Coeruleus Noradrenergic Neurons in Rats Performing T-Maze Tasks.” Scientific Reports 9 (1): 1–13. https://doi.org/10.1038/s41598-018-37227-w.

Yüzgeç, Özge, Mario Prsa, Robert Zimmermann, and Daniel Huber. 2018. “Pupil Size Coupling to Cortical States Protects the Stability of Deep Sleep via Parasympathetic Modulation.” Current Biology. https://doi.org/10.1016/j.cub.2017.12.049.

Zénon, Alexandre, Mariam Sidibé, and Etienne Olivier. 2014. “Pupil Size Variations Correlate with Physical Effort Perception.” Frontiers in Behavioral Neuroscience 8 (August): 1–8. https://doi.org/10.3389/fnbeh.2014.00286.

